# Machine learning applications to predict functional non-coding regions in non-model species

**DOI:** 10.64898/2026.11.27.767885

**Authors:** Jessica A. Peers, Will J. Nash, Samuel A. Speak, Michaël X. Dong, Heather R. Sibley, Kerstin Lindblad-Toh, Ellie E. Armstrong, Klaus-Peter Koepfli, Wilfried Haerty

**Author notes:** Corresponding authors: Jessica A. Peers and Wilfried Haerty. Email addresses.

## Abstract

**Background:** The non-coding genome harbours a wealth of variation associated with diseases and key adaptive traits, but because non-coding variation is difficult to annotate, it is often overlooked in non-model species. Experimental identification of functional non-coding elements, such as promoters and enhancers, is costly and difficult to scale to wild populations. Meanwhile, comparative genomics approaches rely strictly on sequence conservation, which may overlook species-specific patterns. Machine learning (ML) techniques, such as convolutional neural networks, show potential for annotating the non-coding genome, but using such models to make predictions in non-model species has not been thoroughly tested.

**Results:** Here, we use publicly available data from model species to train, evaluate, and benchmark multiple models to predict genome-wide functional non-coding regions, comparing their performance within and across species. We find comparable precision–recall for cross-species predictions of functional non-coding regions. We highlight the issue of class imbalance in the non-coding genome, demonstrating that sequence conservation and transcription factor binding motifs can prioritise true positive results. We apply this approach to the cheetah (*Acinonyx jubatus*), a species with a long-term low effective population size, to identify putatively deleterious non-coding variants across wild and *ex situ* cheetah populations, including variants potentially associated with fertility-related genes previously shown to harbour deleterious coding variants.

**Conclusions:** Our findings show the potential of transfer learning to annotate non-coding genomes in non-model species without species-specific experimental data. We identify class imbalance as a key priority for developing future cross-species ML models.

## Background

Approximately 8-11% of mammalian genomes demonstrate evidence of evolutionary constraint as a consequence of purifying selection acting on functionally important sequences; approximately 80% of these constrained sequences lie outside of protein-coding exons [1,2]. Comparative genomics has enabled the identification of constrained non-coding sequences [1,3,4] and candidate regulatory elements [5], but the functional impacts of variants within these sequences remain harder to interpret than those within protein-coding genes. This difference arises partly because coding sequences are far more tractable experimentally. High-throughput approaches such as cDNA screening and gene expression profiling [6,7] can be readily paired with knockout experiments to test phenotypic effects of specific gene disruptions [8,9] and with saturation mutagenesis, which tests the functional effect of all possible variants in a target region [10–12].

This gap between coding and non-coding annotations persists despite growing evidence for the functional importance of non-coding variation; the vast majority of trait-and disease-associated GWAS hits are found in the non-coding genome, including functional variants implicated in human cancers and autoimmune diseases [13]. This observation extends across model mammal species: a structural non-coding variant causes polled intersex syndrome in goats (*Capra hircus*) [14], and non-coding variants associated with heritable traits, such as brain size and vocal learning ability, have been identified across diverse mammalian lineages [15,16]. Further examples include forelimb development in bats (Chiroptera), associated with regulatory regions located near the *HoxD* gene [17], and a tandem repeat variant in the *DGAT1* promoter region that affects milk fat percentage in dairy cattle (*Bos taurus*) [18].

Because non-coding regulatory elements are difficult to identify and functionally validate, studies of regulatory genome evolution have focused largely on humans and a small number of model organisms [5,19]. Although this focus has yielded valuable insights, the evolution of the non-coding genome in non-model species remains poorly understood. This gap is notable given the enormous phenotypic diversity of mammals, many of whose lineage-specific adaptations have an unknown genetic basis [1]. Functional regions of the non-coding genome are typically annotated using either experimental or comparative genomic approaches, but both have limitations that are especially acute in non-model species.

## Approaches for annotating functional non-coding regions

### Experimental approaches

Functional non-coding regions can be empirically identified using molecular signatures associated with regulatory activity such as chromatin state, methylation status, transcription factor (TF) binding, and histone modifications. Experimental approaches to quantify such signatures include ATAC-seq, DNase-seq, ChIP-seq, Promoter Capture Hi-C, FAIRE-seq and CUT&Tag [20–25]. These techniques can be used both individually and in combination to identify a range of regulatory regions [26].

The data generated by such assays have the potential to be highly accurate but are intrinsically species-specific, and the assays are costly and difficult to apply to non-model species, as high-quality tissue samples are required for most protocols [27–29]. Additionally, as gene expression varies between tissues and developmental stages, representative sampling across tissues and development is essential to develop a comprehensive overview of gene regulation [27,30]; however, sourcing fresh tissues in non-model species can be prohibitively difficult and costly [28].

### Comparative approaches

Comparative genomic approaches offer an alternative option for non-coding annotation, primarily based on the theory that functional regions in the non-coding genome are highly conserved due to purifying selection, showing little change in base identity over evolutionary time [1,31]. Methods such as GERP (Genomic Evolutionary Rate Profiling), PhyloP and PhastCons have been used to identify genome-wide distributions of highly conserved non-coding regions [32–34]. However, GERP has notable limitations: high turnover of functional sequences and non-constant selection can limit detection of loci under selection. Furthermore, Huber et al. [32] identified approximately 80 Mb of human functional non-coding sequence with low GERP scores that would be missed using standard thresholds, suggesting limited sensitivity. PhyloP also has limitations: its resolution varies depending on the number of species compared, from several base pairs using 29 genomes [35] to single nucleotide resolution by integrating over 200 mammalian genomes [1,31], meaning accurate PhyloP scores require large volumes of high-quality reference genomes.

These methods rely on the assumption that sequence conservation directly correlates with functional conservation, but regulatory activity can be maintained despite high sequence turnover [2,36,37]. A further assumption is made that mutations in highly conserved sites are deleterious [32], but sequence conservation alone does not indicate whether a site is functional [38]. Relying on signatures of purifying selection as a proxy for sequence functionality is therefore not always accurate; identified elements may have lost or changed function in the species of interest, species-specific sequences may be missed, and tissue- or cell-type-specific inference is also not possible, as these methods do not incorporate such data.

### Machine learning approaches for functional non-coding annotation

With the increasing availability of taxonomically diverse genomic resources [39–41], there is growing potential for the annotation of constrained genomic regions. Recent advances in machine learning (ML) methodologies enable the incorporation of experimentally validated data from well-studied organisms into the prediction of functional non-coding regions in non-model species. As there is a wealth of experimental data already publicly available for model species, such as humans (*Homo sapiens*) and mice (*Mus musculus*) [42], this integration overcomes limitations associated with experimental methods, such as the lack of relevant species-specific biomaterial. Additionally, ML algorithms can make predictions based on more than just sequence conservation, including sequence features such as transcription factor (TF) motifs and local sequence context, together with experimental data such as chromatin accessibility generated via ATAC-seq. When trained on experimentally validated data, ML predictions can therefore be more accurate than conservation-based approaches like PhyloP [43].

ML methods for the prediction of functional non-coding regions can use several different model architectures, including Support Vector Machines (SVMs), Convolutional Neural Networks (CNNs), Recurrent Neural Networks (RNNs), Bidirectional Long Short-Term Memory networks (BLSTMs) and Bidirectional Encoder Representations from Transformers (BERTs) [44–47]. Since the advent of foundational tools to predict functional non-coding regions, such as gkm-SVM [48], DanQ [49] and Basset [50], CNN-based models have been increasingly used to predict regulatory activity directly from DNA sequence. Basenji [51], the successor to Basset, predicts promoters and distal regulatory elements based on sequences and was successfully transferred between humans and mice. More recent CNN-based tools, such as Puffin [52], DeepEnhancer [53], AI-TAC [54], and ChromBPNet [55], extend these methods to different regulatory prediction tasks. AI-TAC uses deep convolutional neural networks to infer cell-type-specific chromatin accessibility and transcription factor hierarchies from mouse ATAC-seq data. When transferred to human DNA, AI-TAC generated a very similar TF ranking, further highlighting the potential of CNNs for cross-species regulatory inference. Additional ML tools to transfer genomic annotations between species include DeepMEL [56] and TACIT [16].

Using concepts from both CNNs and linear models, ExplaiNN [57] utilises neural additive models, which combine the accuracy of deep learning models with the transparency and interpretability of linear models. Therefore, the benefit of ExplaiNN over other tools is its explainability: the features and properties that contribute to the predictions of the model are clearly provided. This interpretability makes ExplaiNN a good candidate for application to non-model organisms, where transparency of the model’s predictions is particularly important as there is no ground truth to validate predictions against. Additionally, ExplaiNN is provided as a plug-and-play platform, making it accessible to researchers less familiar with ML methods. However, ExplaiNN’s transferability between model and non-model organisms has not yet been tested.

### Non-model species

A broader understanding of the functional impacts of non-coding variation in non-model species is important for both our general understanding of evolutionary biology and for informing conservation strategies, where overlooked functional variation in non-coding regions may be crucial to species survival and persistence [58]. In threatened species, understanding the non-coding genome is particularly important because it can be used to identify deleterious mutations and inbreeding-related genetic load that would otherwise go undetected. Declining populations often experience genetic bottlenecks, leading to the accumulation of deleterious mutations, inbreeding depression, and phenotypic abnormalities [59,60]. Standard population genomics methods, such as calculating runs of homozygosity [61] or identifying deleterious mutations and genetic load in protein-coding genes [62,63], can measure the impact and extent of inbreeding depression. However, without corresponding annotation of the non-coding genome, such methods lack the power to contextualise non-coding variants to understand their functional implications. This is important because mutations in non-coding regions may be similarly deleterious to nonsense or missense mutations in protein-coding exons, due to their potential impact on gene expression [64,65].

The cheetah (*Acinonyx jubatus*) is a non-model species which experienced severe demographic decline approximately 10,000 years ago, resulting in a low effective population size (*N_e_*) [66–68] and inbreeding, with high homozygosity and low nucleotide diversity observed in both wild and *ex situ* populations [66,69–72]. A range of symptoms associated with inbreeding depression have been observed in wild and *ex situ* cheetahs, including poor fertility and morphological and skeletal defects [70,73–76]. Previous studies have identified a range of deleterious variants in protein-coding genes, with an over-representation in male fertility-related genes [66,69,77,78]. However, fully evaluating genetic load and inbreeding depression in the cheetah requires looking beyond the protein-coding regions. Characterizing the non-coding genome will be crucial to untangling the long-term impacts of historical bottlenecks and informing future conservation management.

Here, we test the application of ExplaiNN to predict functional non-coding regions across species, with the aim of annotating the non-coding genome of a non-model species, the cheetah. Using publicly available data from three model species (human, mouse and domestic dog, *Canis lupus familiaris*), multiple ExplaiNN models are trained and tested to determine model performance when transferring between model species. Subsequently, we train a model using input data from all three model species and annotate the functional non-coding genome of the cheetah. We compare model predictions to regions identified by traditional comparative genomics methods and discuss the accuracy of ML compared to existing methods to annotate functional non-coding regions in non-model species. Finally, we intersect our model predictions with a whole genome resequencing dataset to predict variants occurring in functional non-coding regions across *ex situ* and wild cheetah populations.

## Results

### Processing ATAC-seq data and calling peaks

ATAC-seq data from dog, human and mouse were processed using the nf-core atacseq pipeline [79,80] (see Supplementary Methods and Figures S1-S12). Per-sequence GC content showed a roughly normal distribution, suggesting a normal random library in dog, mouse and approximately half of the human samples. Data quality varied by species. Dog samples had lower raw mapping rates (74.4-91.7%) than human (93.3-99.8%) or mouse (96.0-99.1%) samples, as well as high duplication rates (18.3-64.2%). After quality filtering, only 6.6-10.4% of input reads were retained, and far fewer narrow peaks were called per sample in dog (623-5,084) than in human (37,992-340,687) or mouse (17,603-70,592). This loss of reads may reflect the high level of duplication observed in the dog samples. It may also reflect the absence of a mitochondrial genome sequence in the dog reference genome; mitochondrial reads are typically abundant in ATAC-seq libraries [81], and without a mitochondrial contig to map to, these reads would be unmapped and therefore filtered out.

### ExplaiNN: testing on mouse

The first ExplaiNN model generated (Model 1; Table 1) was trained on the mouse, with ‘positive’ sequences defined as 201 bp sequences corresponding to extended ATAC-seq peak summits and ‘negative’ sequences defined as dinucleotide-shuffled positives. For each model, chromosome 1 was removed to act as an independent test set, and the remaining data were split into training, validation and test sets (80/10/10%, respectively). Model 1 was evaluated using performance metrics calculated on the test set: the area under receiver operating characteristic curve (AUC-ROC) = 0.92 and the area under the precision-recall curve (AUC-PR) = 0.92. Model 2 was trained on a similar set of positives and genomic windows with no overlap to ATAC-seq peaks (negatives). When tested on the test dataset, AUC-ROC = 0.90 and AUC-PR = 0.90. Model 2 was then run on the independent test data derived from chromosome 1 and scored AUC-ROC = 0.90 and AUC-PR = 0.90.

**Table 1.** ExplaiNN models trained in this study.

| Model | Species trained on | Positives | Negatives |
|---|---|---|---|
| 1 | Mouse | ATAC-seq peaks extended to 201 bp | Dinucleotide-shuffled Positive sequences |
| 2 | Mouse | ATAC-seq peaks extended to 201 bp | 201 bp genomic windows with no overlap to ATAC-seq peaks or annotated features (repeats hard-masked) |
| 3 | Human |  |  |
| 4 | Dog |  |  |
| 5 | Human and mouse |  |  |
| 6 | Human, mouse and dog |  |  |

Positive sequences were ATAC-seq peaks extended to 201 bp. Negative sequences were either generated by dinucleotide shuffling of the positive sequences (Model 1) or defined as 201 bp genomic windows with no overlap to ATAC-seq peaks or annotated features, with repetitive sequences hard-masked (Models 2-6).

### ExplaiNN: transferring between species

Models 2, 3 and 4 were trained for each species on all chromosomes except chromosome 1. Each model was then tested on chromosome 1 of all three species. Additionally, a multi-species model (Model 5) was trained on combined human and mouse data and tested on combined human and mouse data and on dog data. AUC-ROC and AUC-PR were calculated for each run to evaluate model performance (Figure 1).

**Figure 1.**
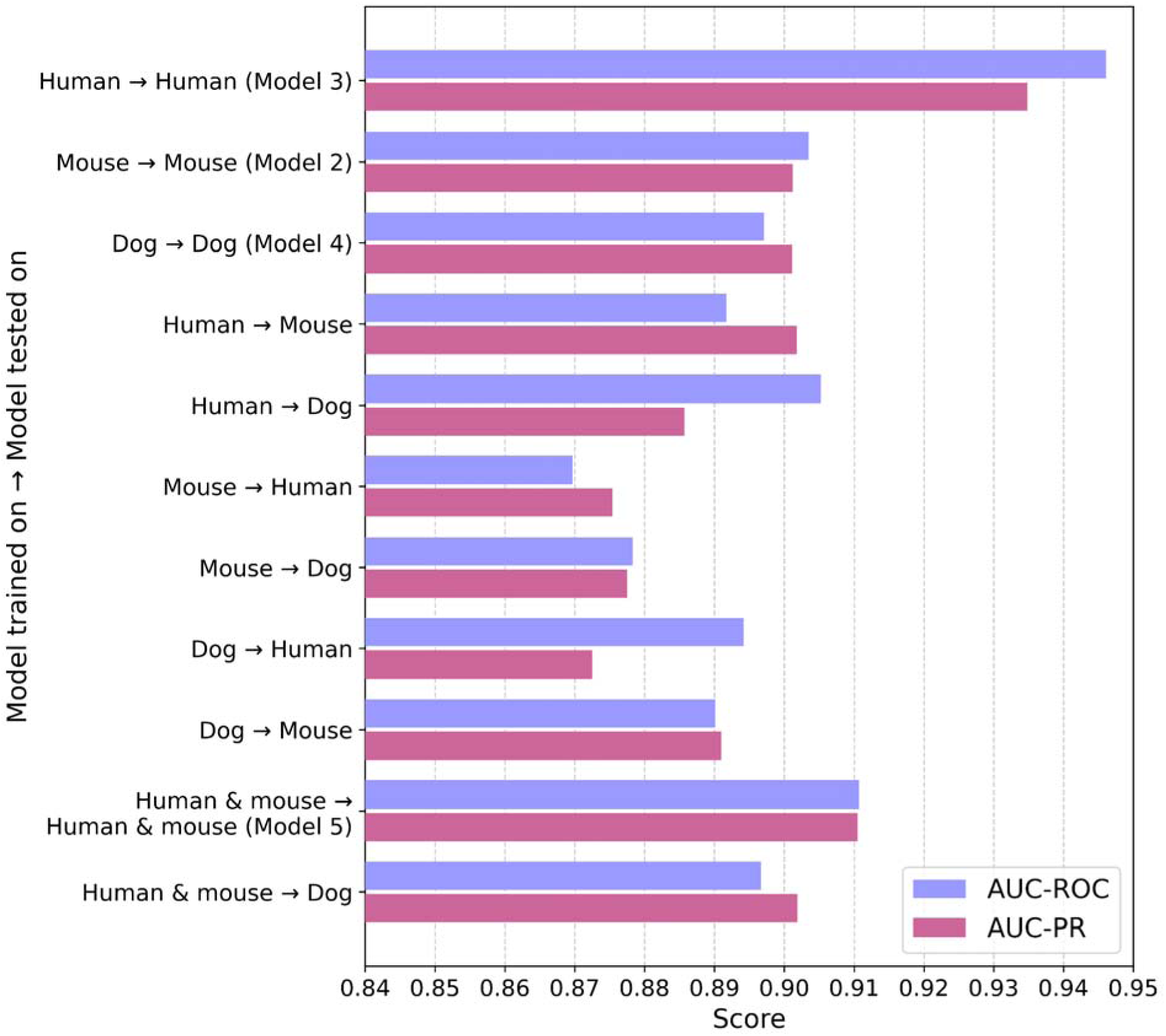
ExplaiNN model performance for within- and between-species non-coding sequence prediction evaluated using AUC-ROC and AUC-PR. Each bar represents the performance of a model trained on chromosomes 2+ from one species (or a combination of species) and tested on chromosome 1 of another species. Model numbers in brackets correspond to Table 1. Area under the receiver operating characteristic curve (AUC-ROC, blue) and area under the precision-recall curve (AUC-PR, pink) are displayed for each model. Intra-species predictions (e.g. human → human) generally outperform cross-species predictions (e.g. human → mouse), but combined training (e.g. human & mouse → dog) improves generalization compared to single-species training.

When ExplaiNN was applied to genome-wide windows rather than balanced datasets, AUC-PR fell markedly in all three species (human: 0.5088; mouse: 0.1875; dog: 0.1368), while AUC-ROC remained relatively high (0.8692, 0.8410 and 0.8096, respectively). This reflects class imbalance rather than poorer model performance: ATAC-seq peaks are rare relative to the number of windows generated across the whole genome (e.g. 14,738 positive versus 1,088,657 negative windows in dog), so even a model with a low false-positive rate produces many more false positives than true positives. AUC-PR is particularly sensitive to this imbalance as it depends on the proportion of positives, so it is not comparable between balanced and imbalanced datasets. Consistent with this, even at very high ExplaiNN prediction scores, predicted windows in the dog were dominated by negatives (Figure 2), and a similar effect was seen in the human and the mouse. Precision, recall and F1 score (the harmonic mean of precision and recall), which are better suited to imbalanced data, were calculated and the number of true and false positives and negatives was quantified for each model (Models 2, 3 and 4; Table 2).

**Figure 2.**
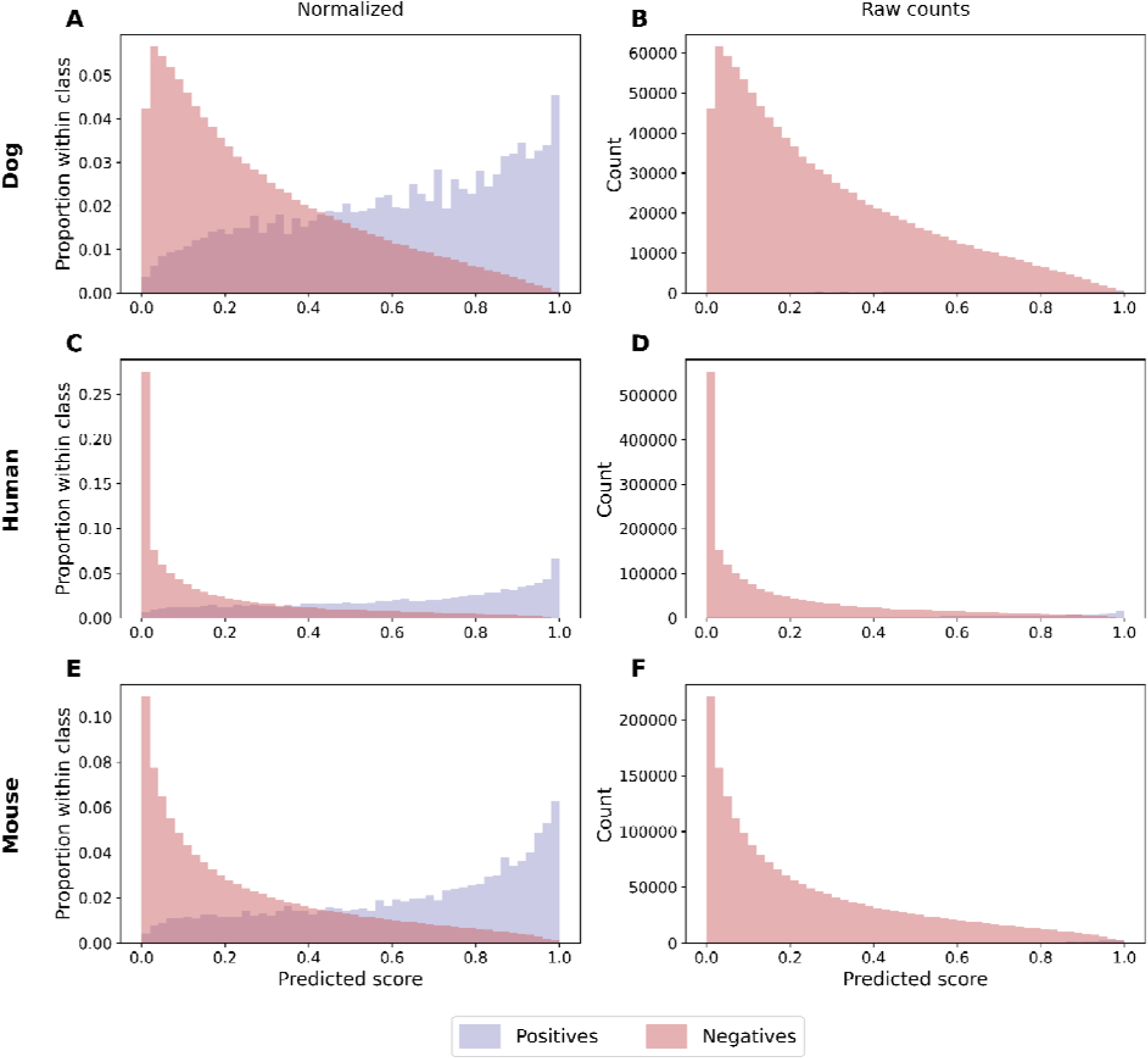
Distribution of ExplaiNN model-predicted scores for 201 bp windows across chromosome 1 of the dog (A & B; Model 4), human (C & D; Model 3) and mouse (E & F; Model 2) genomes. Models were trained on ‘positives’ (ATAC-seq peaks extended to 201 bp) and ‘negatives’ (201 bp genomic windows with no overlap to ATAC-seq peaks or annotated features [repeats hard-masked]) on each species and run on genome-wide windows from each species. **A, C, E:** Normalized distribution of predicted scores, showing the proportion of windows within each class (positive or negative). This view corrects for the class imbalance between positives and negatives, revealing clearer separation in predicted score profiles. **B, D, F:** Raw count distribution of prediction scores without normalization, highlighting the large number of negative windows relative to positives.

**Table 2.** Precision, recall, F1 score, and confusion matrix values for ExplaiNN predictions on genome-wide 201 bp windows.

| **Species** | **Precision** | **Recall** | **F1** | **TP** | **FP** | **TN** | **FN** |
|---|---|---|---|---|---|---|---|
| Dog | 0.04 | 0.65 | 0.08 | 5910 | 206898 | 881746 | 4190 |
| Human | 0.37 | 0.67 | 0.47 | 153220 | 264240 | 1744763 | 76850 |
| Mouse | 0.07 | 0.69 | 0.13 | 26757 | 362221 | 1663723 | 11950 |

To better evaluate model performance under extreme class imbalance (see Figure 2), standard AUC metrics were supplemented with precision, recall, and F1 scores. Results are shown for models trained and tested on each species (dog, human, mouse). True positives (TP), false positives (FP), true negatives (TN), and false negatives (FN) are also reported.

### ExplaiNN: reducing number of false positives

Due to the difficulty of discerning true and false positives, ExplaiNN prediction scores, conservation scores, and distance to nearest coding region were used to filter false positives in the dog. Prior to any filtering of predictions, ExplaiNN labelled 5,910 true positives and 206,898 false positives – a ratio of 1:35 (Table 2). To account for the overrepresentation of false positives introduced by class imbalance, we examined the distribution of ExplaiNN prediction scores across the four categories (TP, FP, TN, FN). Based on pairwise Mann–Whitney U testing, significant differences (p < 0.05) between the distributions of ExplaiNN scores for TP and FP and for TN and FN were identified, as well as all other combinations of pairs of categories (Figure 3A). When filtering the set of TP and FP for ExplaiNN prediction scores ≥ 0.75, 3,162 TP and 54,038 FP remained for the dog, a ratio of 1:17.

**Figure 3.**
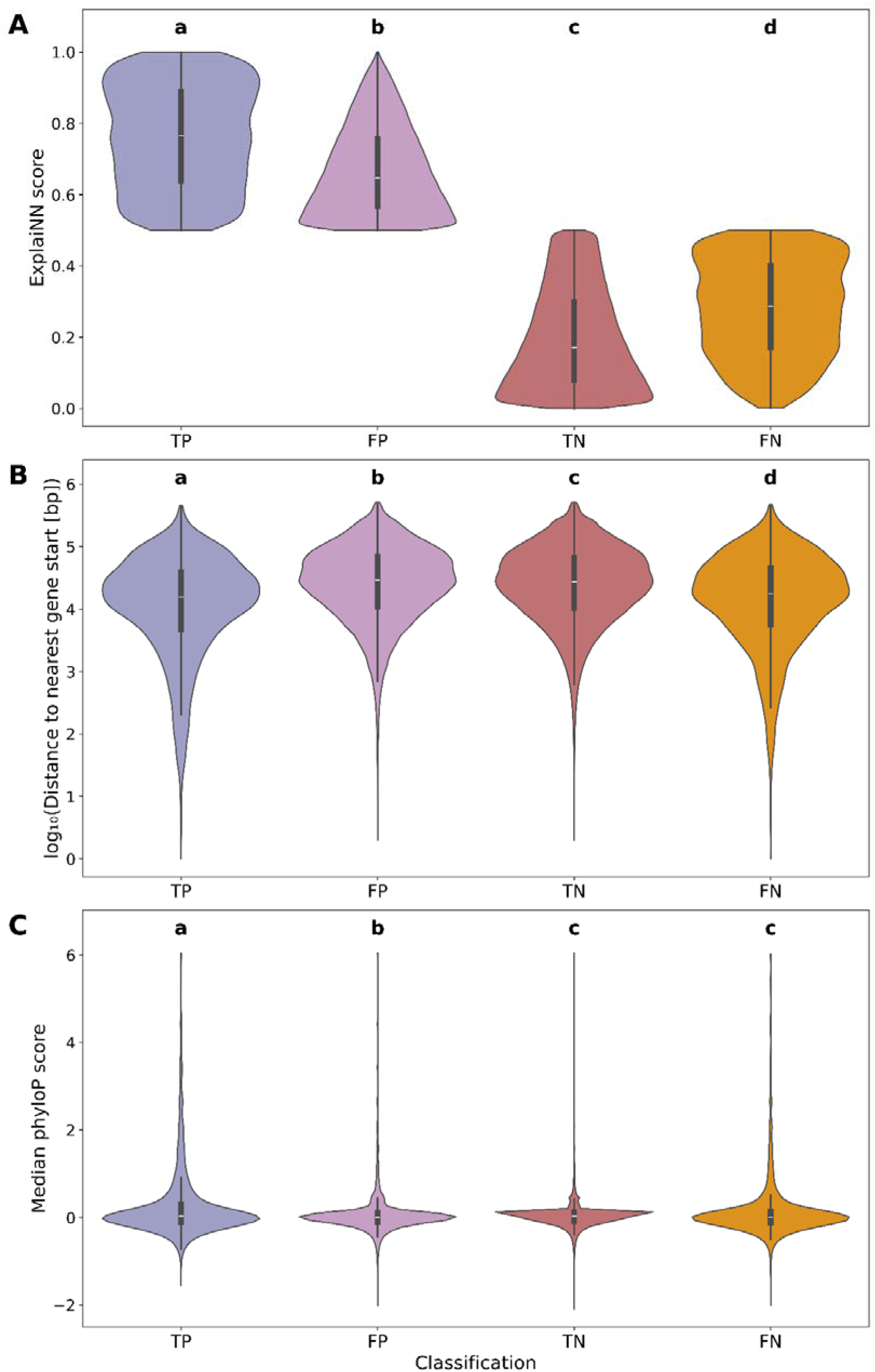
Features associated with predictions in the domestic dog. (**A**) **Distribution of ExplaiNN prediction scores by classification category.** The distribution of prediction scores for each of the four classes: true positives (TP), false positives (FP), true negatives (TN), and false negatives (FN). All pairwise comparisons showed significance (p<0.001, indicated by a, b, c, d annotations). (**B**) **Log-transformed distance to the nearest downstream gene for each classification category.** The distribution of distances (log_10_-transformed) from the end of each 201 bp window to the nearest downstream gene, stratified by classification outcome. Statistical significance was assessed using the Mann–Whitney U test, with all pairwise comparisons showing significance (p<0.001, indicated by a, b, c, d annotations). (**C**) **Median PhyloP score per window across classification categories.** The distribution of median PhyloP conservation scores for each classification type. Mann–Whitney U tests indicated significant differences between most category pairs (p<0.001), with the exception of TN vs FN (p = 0.193), as indicated by labels a, b, c. See Table S5 for full statistical outputs.

When comparing distance to gene start site for each of the categories, a significant difference (Mann–Whitney U, p < 0.05) was found between all pairs of categories (TP, FP, TN, FN; Figure 3B). When filtering the raw set of TP and FP for a distance to the nearest gene start site of ≤ 50,000 bp, 4,723 TP and 136,203 FP were observed, a ratio of 1:29. When comparing median PhyloP score for each category (Figure 3C), a statistically significant difference was found between true and false positives (Mann–Whitney U, p<0.05) but not between true and false negatives. When filtering by PhyloP score ≥ 2, the set of true and false positives was 277 TP and 2,824 FP, resulting in a ratio of 1:10. When all three filters were run sequentially, the final set of true and false positives was 123 TP and 551 FP, resulting in a ratio of 1:4.5.

Finally, we examined which TF binding motifs the ExplaiNN models had learned. ExplaiNN filters, 19-bp sequence motifs that contribute to the model’s predictions, were extracted and compared with known TF motifs in the JASPAR 2024 Vertebrate Core Non-redundant motif database. For the dog, of the 100 most informative filters, only 15 matched known TF motifs (Table S6). For human and mouse, 17 and 19 filters matched known TF motifs, respectively. The importance of each of these TF-matching filters to model predictions was plotted per species (Figure 4A, Table S6).

**Figure 4.**
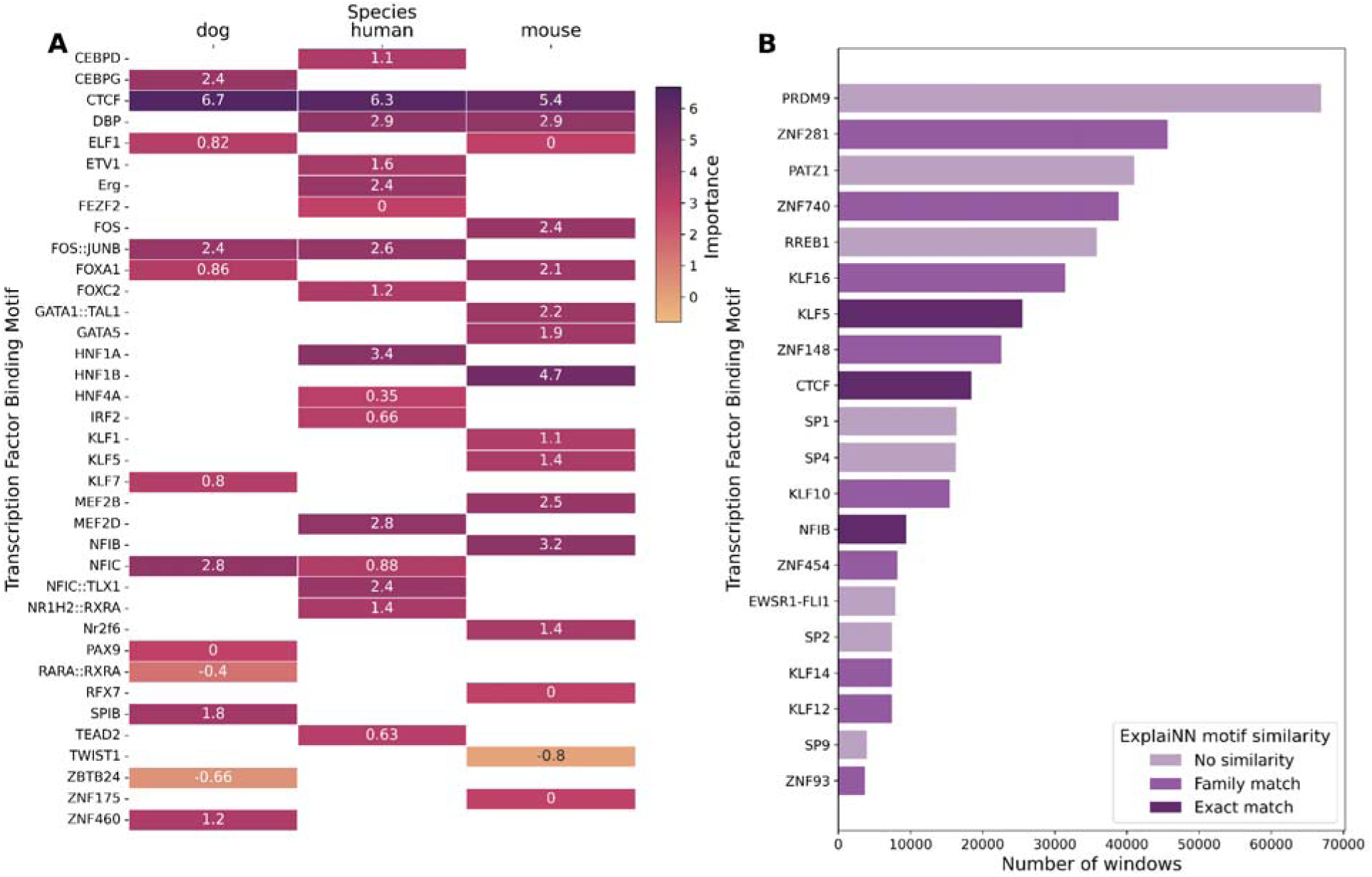
ExplaiNN predicts windows containing a range of transcription factor (TF) binding motifs. (A) Motif importance by species for ExplaiNN predictions. The relative importance scores of TF motif-like filters learned by ExplaiNN for dog, human, and mouse models are shown. Each row represents a distinct motif, and each column corresponds to a species. Where a motif filter appeared multiple times within a species, the highest importance score was retained. Higher scores indicate greater contribution to model predictions. White signifies no occurrences of the motif in that species. **(B) Transcription factor binding motifs identified in predicted functional non-coding windows in the cheetah.** The twenty most frequently occurring transcription factor motifs and their number of significant hits (q-value < 0.05, FIMO) across the final set of predicted functional non-coding windows. Colour indicate similarity to motifs used by ExplaiNN for prediction in (A): light purple = no similarity, medium purple = similarity at the protein family level, and dark purple = exact match.

### ExplaiNN: running on cheetah and comparing to sequence conservation

When ExplaiNN Model 6 (trained on human, mouse and dog; Table 1) was applied to the cheetah genome, 10,887,913 windows were predicted to be ‘positive’ (prediction score ≥ 0.5), and 36,659,747 windows were predicted to be ‘negative’ (prediction score < 0.5). The total set of windows was filtered to retain those outside of protein-coding genes, leaving 44,330,717 windows. Windows were then filtered to retain those with ExplaiNN prediction scores ≥ 0.75, leaving 3,194,395 positive predictions. We excluded 6,263 windows located on unplaced contigs/scaffolds due to poor alignment quality, retaining only those on the 19 chromosomal scaffolds, which comprise 99.72% of the reference genome assembly. Remaining windows more than 50,000 bp from the nearest downstream gene start site were filtered out, leaving 1,767,739 windows. Of these, only 542 had a mean PhyloP score ≥ 2, all of which overlapped a UCE. Therefore, the intersection of the 1,767,739 positive ExplaiNN windows and UCEs was extracted, resulting in 292,912 remaining windows. Because filtering for PhyloP score was too stringent for our data, we used UCE-intersect as a ‘sequence conservation’ filter instead, taking this set as the ‘final’ set of predicted functional non-coding regions.

A skewed distribution of distances to the nearest gene start site was observed (Figure S13), with the highest density occurring within approximately 2-5 kb upstream of the gene, and the majority of predicted functional windows located within ∼10-15 kb. Of the 292,912 windows, 94,945 had at least one significant FIMO motif hit. The top 20 most common TF binding motifs predicted were extracted (Figure 4B) and compared to the motifs learned by ExplaiNN in Figure 4A: three were exact matches for ExplaiNN motifs and nine were within the same protein family as ExplaiNN motifs.

### Variant annotation in the cheetah

We implemented a multi-stage filtering scheme (Table 3) to prioritise potentially functional variants from the 4,013,408 whole-genome SNPs identified across 39 cheetahs [69]. Restricting variants to predicted functional non-coding windows retained 77,931 SNPs, of which 28,345 had a CADD or NCBoost score (Figure S14). Applying a 75th-percentile threshold for either metric (CADD > 7.61 or NCBoost > 0.86) yielded 7,384 high-impact candidate SNPs. Finally, intersecting these candidates with significant FIMO motif hits isolated 4,840 putatively deleterious non-coding variants for downstream population genomic analyses. These SNPs overlapped 728 motifs, most frequently CTCF (CCCTC-binding factor), TFAP (Transcription Factor Activating enhancer binding Protein) and ZNF (Zinc Finger Protein) motifs (Table S7).

**Table 3.** Sequential filtering steps used to identify putatively deleterious non-coding variants in cheetah genomes.

| **Stage / Filter Step** | **Filtering Criteria / Threshold** | **SNPs Retained** | **% Retained (Prior Step)** | **% of Total SNPs** |
|---|---|---|---|---|
| **1. Total Resequencing SNPs** | Whole-genome variant calling across 39 cheetahs [69] | 4013408 | 100.0% | 100.0% |
| **2. Functional Non-Coding** | Localized within predicted regulatory windows | 77931 | 1.94% | 1.94% |
| **3. Functionally Scored** | Scored by CADD (n=5766) or NCBoost (n=25288)<sup>*</sup> | 28345 | 36.37% | 0.71% |
| **4. High-Impact Candidates** | Exceeding 75th percentile: CADD > 7.61 or NCBoost > 0.86<sup>**</sup> | 7384 | 26.05% | 0.18% |
| **5. Motif Disruption (Final Set)** | Overlapping significant FIMO TFBS prediction (n=728 motifs) | **4840** | **65.55%** | **0.12%** |

SNPs were filtered by location within predicted functional non-coding windows, CADD and NCBoost scores, and overlap with significant FIMO hits. <sup>\*</sup>A total of 2,709 SNPs received both CADD and NCBoost scores. <sup>\*\*</sup>Comprises 1,442 high-CADD SNPs, 6,326 high-NCBoost SNPs, and 384 SNPs meeting both thresholds.

To associate putatively deleterious SNPs with the genes they may impact, TADs were called from cheetah Hi-C data. Of the 340,869,167 read pairs sequenced, 97.71% aligned to the cheetah reference genome. After removing PCR and optical duplicates, 226,503,637 unique read pairs (66.45%) remained, with an estimated library complexity of 797,749,419. Of these, 207,835,298 read pairs (60.97% of all read pairs) passed the MAPQ ≥ 30 threshold and represented valid Hi-C contacts, comprising 185,206,198 intra-chromosomal (54.33%) and 22,629,100 inter-chromosomal (6.64%) contact pairs. Of intra-chromosomal contacts, 68.31% were short-range (<20 kb) and 13.45% were long-range (>20 kb). Arrowhead identified 5, 106 and 503 TADs at 10 kb, 25 kb and 50 kb resolution respectively, with genome coverage increasing accordingly (0.16%, 7.37% and 25.47%). The low number of TADs is likely caused by the low coverage of the Hi-C data. The 50 kb-resolution TADs were therefore used for all downstream analyses.

Of the 4,840 putatively deleterious non-coding SNPs, 840 (17%) fell within a TAD. Using TADs to associate these SNPs with genes resulted in 519 unique genes (‘TAD-informed’ gene set). Of these, 207 mapped to a human Ensembl ID. When associating genes with SNPs solely based on genomic proximity, 3,173 unique genes (‘proximity-informed’ gene set) were associated with the 4,840 SNPs, of which 1,732 mapped to a human Ensembl ID. In total, 469 genes were identified in both the ‘TAD-informed’ and ‘proximity-informed’ gene sets, resulting in a union of 3,223 unique genes. GO enrichment analysis identified no significant functional enrichment (FDR-corrected q-value > 0.05) in either set of genes or the union of the two sets.

Of the 3,223 total genes associated with the 4,840 high-impact SNPs, 829 had previously been observed to contain deleterious mutations within protein-coding exons (Table S4). Of these, 745 were found in one publication/source, 81 were found in two sources, and 3 genes were found in three sources. GO enrichment analysis found no significant enrichment in any set of genes. The genes found in three sources were IFT140, PANX1, and SHBG, which have previously been observed to contain high- and moderate-impact SNPs classified by SnpEff [69] and deleterious variants (SIFT score 0-0.05) [78].

We examined the distribution of the 4,840 putatively deleterious non-coding SNPs across populations; each population contained at least 88 unique SNPs. The lowest proportion of these deleterious SNPs was observed in Tanzania, with the highest proportion in the U.S. population (Figure 5A&B).

**Figure 5.**
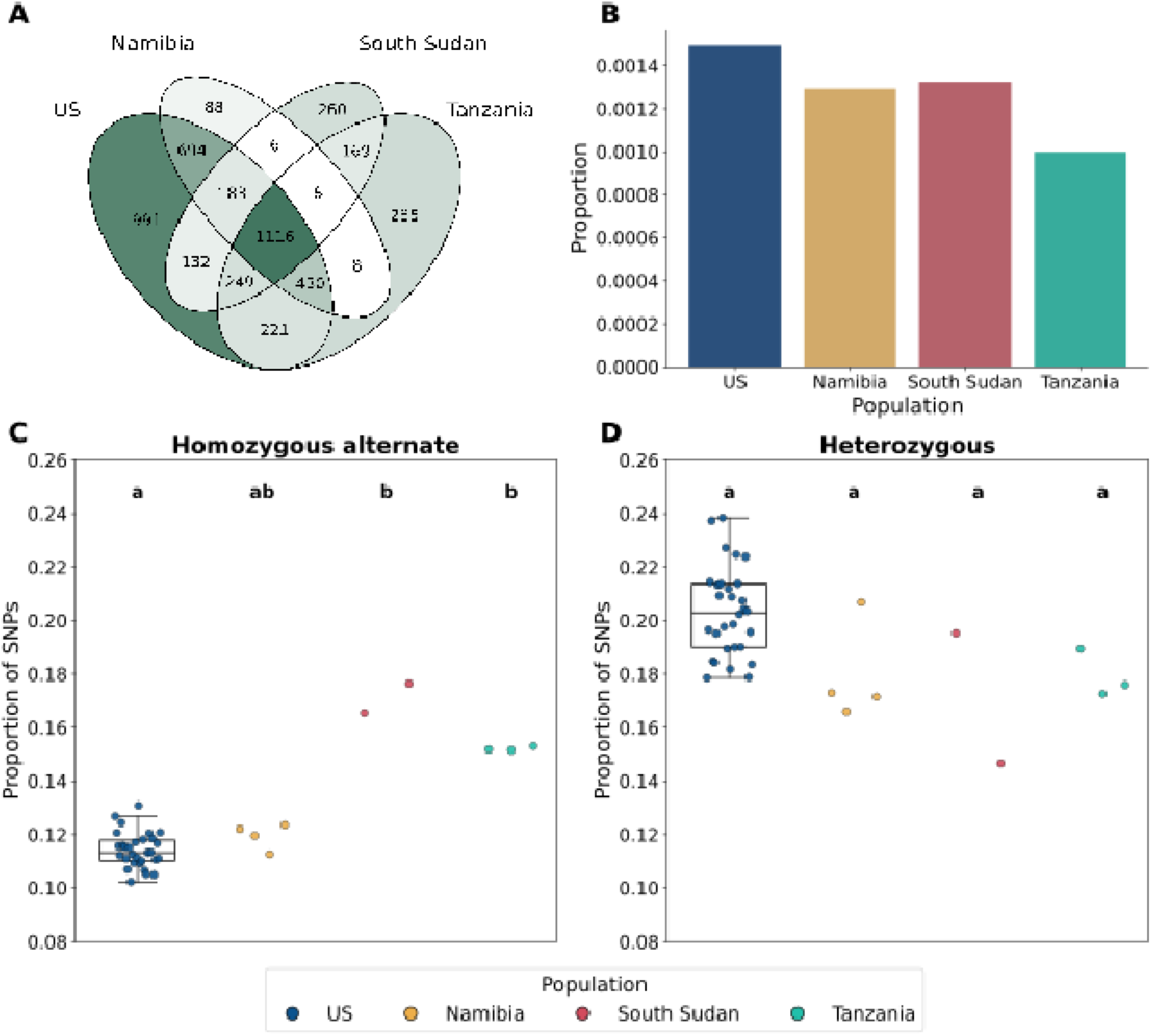
Putatively deleterious non-coding SNPs across wild and *ex situ* cheetah populations. Colours indicate populations: U.S. (blue), Namibia (yellow), South Sudan (red), and Tanzania (green). Compact letter display (a, b) in panels C and D shows the results of pairwise Mann–Whitney U tests with Bonferroni correction (p < 0.05), where populations sharing a letter do not differ significantly. (**A**) Distribution of 4,840 putatively deleterious non-coding SNPs between three wild (Namibia, South Sudan, Tanzania) and one *ex situ* (U.S.) cheetah population. The number of SNPs specific to each combination of populations is shown in each segment, with a darker colour indicating a higher number of SNPs. (**B**) Putatively deleterious non-coding SNPs unique to each population as a proportion of the overall number of unique SNPs per population (see Figure S11 in [69]). (**C**) Proportion of homozygous alternate and (**D**) heterozygous putatively deleterious non-coding SNPs per sample, calculated as a fraction of all putatively deleterious non-coding SNPs with a called genotype in that sample.

The proportion of putatively deleterious non-coding SNPs that were homozygous for the alternate allele or heterozygous was calculated per sample, out of all such SNPs with a called genotype in that sample (Figure 5C&D). A significant overall difference was identified for both the homozygous alternate proportion (Kruskal–Wallis: H = 15.0395, p = 0.00178) and the heterozygous proportion (Kruskal–Wallis: H = 11.5972, p = 0.00890). Bonferroni-corrected pairwise Mann-Whitney U tests identified significant differences in the homozygous alternate proportion between the U.S. and South Sudan and the U.S. and Tanzania (p = 0.0242 and p = 0.00220, respectively), while all other pairwise comparisons were non-significant. The homozygous alternate proportion was highest in South Sudan and Tanzania (median proportion: South Sudan = 0.171, Tanzania = 0.152, Namibia = 0.121, U.S. = 0.113). The heterozygous proportion had the highest median in the U.S. (median: U.S. = 0.203, Tanzania = 0.176, Namibia = 0.172, South Sudan = 0.171), but pairwise Mann–Whitney U tests identified no significant difference between any pairs of populations after Bonferroni correction (p > 0.05).

## Discussion

Here, we present the novel application of an ML tool, ExplaiNN, to predict functional non-coding sequences in a non-model species. Through comparative analysis of chromatin accessibility data sourced from a suite of model mammals, we apply this method to the cheetah. We demonstrate that it is possible to train a model that accurately annotates the non-coding genome, with the caveat that data imbalance, caused by the relatively small amount of regulatory sequence compared to non-regulatory sequence in the non-coding genome, impacts the accuracy of this method. These findings provide support for the utility of this novel approach in studying the non-coding genome of non-model species, expanding the potential of research into crucial trait-and disease-associated variants in such species, and highlight the need for future ML tools to consider the issue of data imbalance.

### Testing ExplaiNN within species

Our initial ExplaiNN models were run on model species to test model performance against a ground truth (ATAC-seq peaks), taking advantage of publicly available ATAC-seq data in the human, mouse and domestic dog. Whilst the majority of human and mouse data showed low read duplication and a high number of reads passing filtering steps, the processing steps highlighted several issues with the dog data, namely high read duplication and low mapping rate, resulting in a large proportion of reads being filtered out. Although larger data volumes are generally considered better for model training [82], models trained on this filtered subset performed as well as those trained on the full dataset, demonstrating that filtering did not impact model performance.

The first ExplaiNN model (Model 1; Table 1) was trained and tested using synthetic negatives (dinucleotide-shuffled positive sequences), raising the question of whether the model learned to distinguish true ATAC-seq peaks from background sequences, or simply real from synthetic DNA. As expected, and consistent with Novakovsky et al.’s [57] observations, performance dropped when the model was instead trained and tested using real negatives, confirming that the initial high performance partly reflected this real-versus-synthetic distinction rather than genuine peak-calling ability. This supported the use of real negatives in all subsequent models.

Our ExplaiNN models were trained on 201 bp sequences as recommended by Novakovsky et al. [57], as ExplaiNN requires all sequences in the training, testing, and predicting sets to be the same length. Individual regulatory elements such as enhancers and promoters typically span ∼50 bp to several kb [83,84]. Therefore, limiting the model to only predict on 201 bp regions may not be as biologically informative, and extending the model to encompass the diversity of lengths of functional non-coding sequences is an important step in any future work.

### Testing ExplaiNN across species

Although ExplaiNN is not necessarily designed for cross-species prediction, it performed well (AUC metrics approximately 0.87–0.91) when trained on a different species from the species it was tested on. Performance (AUC-ROC and AUC-PR) was nevertheless lower than within-species models, consistent with [85], who suggest that cross-species neural networks perform worse than within-species models. In contrast, a model trained on multiple species performed as well as within-species models, highlighting the importance of multi-species training data. We tested model performance across an evolutionary distance of approximately 82.5 million years [86], indicating that with sufficient or multi-species training data, our approach should generalise to other placental mammals. This represents a novel cross-species application of ExplaiNN, allowing non-model species to be studied using only model-species training data.

To form our multi-species training dataset, we chose mouse and human for the quality and volume of their data and domestic dog for its smaller evolutionary distance to the cheetah. We included heart, stomach, and liver, the only tissues available in all three species, to ensure equal representation across species, and combined them into one dataset to reduce tissue specificity effects. The model may therefore predict regulatory elements of these tissues but overlook those of others, meaning some ‘false positives’ may arise from regulatory elements in other tissues that the model recognises as having a similar architecture to those in the tissues we provided. Thorough testing with a wider set of ATAC-seq data is needed to confirm this possibility. It is also possible that ATAC-seq data from more species, particularly closer relatives such as the domestic cat, or more tissues may improve model predictions further. However, as predictions were accurate on balanced datasets, more data is unlikely to reduce the number of false positives. Finally, although we show that ML tools can be applied to species with little experimental data beyond a reference genome, the models are trained on extensive experimental data from other taxa. Their performance may therefore decline when data from closely related species or relevant tissues are absent from the training set.

The primary issue we encountered was data imbalance; when the non-coding genome is broken into windows, there are considerably fewer positive windows (ATAC-seq peaks) than negative windows (non-coding regions with no overlap to ATAC-seq peaks). Although the training and testing data could be artificially balanced to confirm model performance, the mammalian genome is inherently unbalanced, with far fewer regulatory elements than non-functional non-coding sequence [1,2]. Even with very high model performance (AUC metrics > 0.9), a high number of false positives (windows with no ATAC-seq peak overlap that were predicted as positive by ExplaiNN) therefore remain in an imbalanced dataset, as observed in the dog. As this work aimed to annotate functional non-coding regions in a non-model species with no available data or annotation, it is not possible to force the dataset to be balanced. Therefore, additional metrics were required to separate true and false positives.

Firstly, we tested sequence conservation, which existing methods use as a proxy for function, assuming that functional regions are under strong purifying selection and will therefore be conserved between species [1,2,63]. True positives are therefore expected to have higher sequence conservation than false positives. Conservation differed significantly between true and false positives, although there was still considerable overlap and this statistical significance may, in part, reflect the large number of values in the dataset. However, selecting the putative functional non-coding regions with the highest conservation scores removed a higher proportion of false positives than true positives, giving higher confidence in the model predictions. Although sequence conservation does not necessarily indicate function, as some conserved sequences are not functional and some functional sequences are not conserved [87], the number of false positives in the dataset made this filtering step necessary.

As sequence conservation alone did not accurately distinguish between true and false positives, the location of each window relative to the nearest downstream coding region was also considered. Although the distance between a functional non-coding region and a gene can vary, most promoters lie within a few kilobases of coding sequences [88–93]. Therefore, true positive windows were expected to be closer to coding regions than false positive windows. Again, a statistically significant difference was observed, with considerable overlap between true and false positives, but filtering for distance to coding regions improved the true-to-false-positive ratio. This step is affected by the quality and method of genome annotation used for the non-model species. The cheetah genome annotation used here (VMU_Ajub_asm_v1.0; [94]) was generated by a homology-based approach using nine mammalian genomes as references and is therefore likely to lack cheetah-specific orphan genes and complex gene families (e.g. immune genes) which are poorly annotated by automated approaches [95]. Restricting the set of positive predictions to those within 50 kb of annotated genes may therefore miss regulatory elements upstream of unannotated complex gene families, but these gene families typically require extensive manual curation [95], which was beyond the scope of this study.

Another method we used to separate true and false positives was overlaying transcription factor binding motifs onto the windows, as functional regulatory regions are typically enriched for TF binding sites [5]. Among the filters of ExplaiNN (the features generated by the model) that were most important in making predictions, a relatively low proportion matched known TF motifs. Most of these filters had positive importance scores, indicating that they contributed primarily to predicting positive sequences (i.e. ATAC-seq peaks). The motif with the highest importance, CTCF, is critical for gene activation and repression, chromatin looping and enhancer blocking [96–99]. Three motifs had negative importance scores, suggesting that they were more important for predicting negative sequences: RARα–RXRα (a heterodimer key in retinoid signalling; [100]), TWIST1 (involved in embryonic development; [101]) and ZBTB24 (involved in DNA methylation at centromeres; [102]). Given their specialised functions, these motifs are unlikely to be associated with ATAC-seq peaks from the heart, liver and stomach samples used here, and the genes encoding these TFs have low expression in humans [103].

As ExplaiNN’s motif selection is consistent with our understanding of transcriptional regulation, this, combined with the other filtering parameters, gave confidence in our final set of predicted ‘positives’. However, as discussed previously, we still expect this set to contain false positives because of the imbalanced nature of the genome. Although this issue is not unique to ML methods, it particularly affects classification tasks, where the rare true positive class can be swamped by false positives. We therefore compared our predictions to existing methods of predicting functional non-coding regions: TF motif scanning and ultra-conserved elements (UCEs). The vast majority of predicted windows contained at least one significant hit for a TF binding motif, which suggests functionality, although a motif-like sequence does not guarantee a functional motif [104]. Most predicted windows also overlapped a UCE, suggesting a high correlation between ExplaiNN’s predictions and sequence conservation. Windows with no overlap to a UCE may still be functional, as function can be conserved even when sequence is not [2,38]. However, given the lack of species-specific data and the severely imbalanced training and testing data, it is not possible to determine whether these predictions are accurate.

As discussed, the most critical issue with current machine learning tools for functional non-coding prediction is that genomic data are inherently and severely imbalanced [105]. Even with the most accurate classifier trained on a large amount of data, the sparsity of functional non-coding regions throughout the genome means that predictions may suffer from low precision, with many false positives. This issue persisted here even when alternative performance metrics were used, with both precision and recall affected. Class imbalance is a common problem in machine learning, particularly in genomics [106,107], and future machine learning tools must be built with this imbalance in mind to enable robust predictions and downstream biological interpretation. Genomic language models (gLMs), such as GPN-Star [108], DNABERT [109] and Evo 2 [110], learn sequence ‘grammar’ across the whole genome and may offer a partial solution, but they do not remove the underlying imbalance of the genome. In addition, although some such models (e.g. NTv3, PlantCaduceus, and NucEL [111–113]) have been tested across species, to our knowledge none has been evaluated on non-model mammals.

### Variant annotation in the cheetah

When applying ExplaiNN to the cheetah, we identified variants within predicted functional non-coding windows. Despite several filtering steps, class imbalance means that some of these predicted functional windows are likely to be false positives. We therefore further filtered variants within these windows by CADD score, NCBoost score, and overlap with a predicted TFBS to prioritise putatively deleterious variants. However, each of these filters has caveats. Firstly, both CADD and NCBoost scores were lifted over from the human, assuming that a variant that is deleterious at a given position in the human is similarly deleterious in the cheetah. Species-specific genome features may make this untrue, particularly for NCBoost scores outside UCEs, where sequence conservation is weaker and species-specific differences are more likely to affect accuracy. Secondly, the 75th-percentile thresholds we applied to CADD and NCBoost scores are arbitrary and may exclude deleterious variants below the threshold or include non-deleterious variants above it. Finally, a TF motif-like sequence does not guarantee a TFBS, so some variants may not disrupt a binding site. Without cheetah-specific data, we cannot verify these assumptions, but these filters allowed us to prioritise the non-coding variants that were most likely to be deleterious.

Several methods were used to associate predicted deleterious variants with genes, including taking the nearest downstream gene and considering genes within the same TAD, although the latter was limited by low-coverage Hi-C data which resulted in a low proportion of the genome being called in TADs. However, gene regulation is non-linear, so the variants we identify do not necessarily affect the genes in closest proximity [114,115]. Additionally, this functional annotation relies on multiple layers of prediction (predicted functional non-coding window, predicted deleterious impact of each variant, predicted gene associated with each variant), each with the potential for false positive results, so these variant annotations should be interpreted with caution. For more accurate association between non-coding regions or variants and genes, recent methods such as SCENT, Signac, and pgBoost [116–118] integrate genomic distance with multi-omic data to predict SNP-gene links, but to our knowledge, these methods have not yet been tested in non-model species and require significant multi-omic data, which were not available for this study.

Although no significant enrichment for gene function was observed in genes associated with putatively deleterious SNPs in functional non-coding regions, we found 829 genes that have previously been associated with deleterious mutations in protein-coding exons [66,69,77,78,119]. These are therefore candidate genes that warrant further investigation as possible contributors to inbreeding depression phenotypes. Three genes were found in three sources: *IFT140* (Intraflagellar transport protein 140), *PANX1* (Pannexin 1) and *SHBG* (Sex hormone-binding globulin). *IFT140* is involved in ciliary/flagellar assembly and is essential for male fertility and spermiogenesis in mice [120], *PANX1* is involved in oocyte function and female fertility [121], and *SHBG* is involved in regulating androgens and oestrogens [122]. These genes all have roles in male and/or female reproduction, as expected given that previous studies focused on reproduction-related genes, and could be important candidate genes for future work to uncover the causative mutations underlying poor fertility in cheetahs [123]. However, further work to confirm the deleteriousness of the variants we identify and their association with these genes is necessary.

We also found that the distribution of deleterious non-coding SNPs mirrors that of the coding SNPs [69], with a lower proportion of population-specific SNPs in Tanzania. The proportion of homozygous alternate SNPs was significantly higher in South Sudanese and Tanzanian cheetahs compared to the U.S., again mirroring the pattern observed in coding SNPs [69]. Although strong evidence of inbreeding in the Tanzanian population was previously observed, this is not reflected in the number of population-specific deleterious SNPs. This result highlights the importance of considering non-coding mutation load in conservation genomics, as each population contains unique deleterious non-coding variation which could be contributing to inbreeding depression phenotypes and decreased fitness. Future work should validate the mutations we identify here to confirm their functional impact, enabling their use in informing conservation management in both wild and *ex situ* populations. Additionally, sample sizes for the wild populations (Namibia, South Sudan, and Tanzania) are very low, meaning the patterns we observe in SNP distribution and load may be skewed by this sampling; additional samples are required to confirm our observations and accurately inform conservation management. It is also important to consider the effects of reference bias. The reference genome we used was generated from an *ex situ* cheetah in Lisbon Zoo, Portugal, which does not belong to any of the four populations we sampled. However, this individual may be most closely related to the U.S. *ex situ* population or the Namibian population, as the southern African subspecies is the likely source population of most *ex situ* cheetahs [124]. The number of deleterious variants in these populations may therefore be underestimated. Additionally, any deleterious non-coding mutations fixed across the species, or at least across our samples, are not detected because they do not differ from the reference, so further deleterious non-coding mutations may remain to be identified.

Species-specific functional genomic data would address several of the limitations noted above. Training or fine-tuning interpretable models such as ExplaiNN [57] on data from the target species would better capture its tissue-and cell-type-specific regulatory landscape and provide a ground truth for evaluating predictions, which we lacked here. Although such data would not remove the underlying class imbalance of the genome, they would enable more accurate annotation of putatively deleterious non-coding variants and variant-gene links. Building on the Functional Annotation of Animal Genomes (FAANG) framework (Andersson et al. 2015), we suggest generating chromatin accessibility (ATAC-seq), histone modification (ChIP-seq or CUT&Tag), and gene expression (bulk or single-cell RNA-seq) data from tissues spanning major organ systems. We recognise that generating these data is costly and constrained for non-model species, particularly those classed as endangered, where permits and limited access often restrict sampling. However, species managed in *ex situ* breeding programmes, such as the cheetah and black-footed ferret (*Mustela nigripes*), are notable exceptions because tissues and viable cell lines are routinely banked for research [125]. Expanding biobanking beyond material collected for reference genomes to include a broader range of tissues and viable cells would therefore support the building of functional genomic resources in non-model species that are comparable in scope to those available for human and mouse [19].

## Conclusion

In this study, we show that ExplaiNN, a CNN-based machine learning method, can predict functional non-coding regions across species, with cross-species models performing comparably to within-species models. We applied this approach to the cheetah to annotate its non-coding genome and identify putatively deleterious non-coding variants, including variants potentially associated with fertility-related genes. However, class imbalance in genomic datasets introduces substantial noise in the form of false positives. Additional filters, including sequence conservation, CADD and NCBoost scores, and TF motifs, only partly mitigate this, so our predictions should be interpreted with caution. Our results demonstrate the potential for machine learning methods to predict functional non-coding regions in non-model species, but further methodological development to handle class imbalance is needed before such predictions can be used with confidence. Nevertheless, our framework offers a novel approach to annotating non-coding genomes of non-model species without the requirement for species-specific experimental data.

## Methods

### Genomic and ATAC-seq data

Publicly available ATAC-seq data for human, mouse and domestic dog were used in this study. For the human and mouse, raw ATAC-seq read files were downloaded from ENCODE (https://www.encodeproject.org/) (Table S1) [5]. Dog ATAC-seq read files were downloaded from BarkBase (https://www.barkbase.org/) (Table S2) [126]. To establish a balanced multi-species training dataset and mitigate species-specific bias, we included tissue types shared across all three species: stomach, heart and liver. For human, mouse and domestic dog, 29, 18 and 10 samples were downloaded, respectively (Table S1 and S2). The corresponding reference genome for each species was downloaded from NCBI (Table 4).

**Table 4.** Reference genomes used for ATAC-seq read processing. Scientific and common names are provided for each species alongside genome assembly and NCBI RefSeq accession

| Species | Common name | Genome | NCBI RefSeq |
|---|---|---|---|
| *Canis lupus familiaris* | Dog | ROS_Cfam_1.0 | GCF_014441545.1 |
| *Mus musculus* | Mouse | GRCm39 | GCF_000001635.27 |
| *Homo sapiens* | Human | GRCh38.p14 | GCF_000001405.40 |

### Processing ATAC-seq data and calling peaks

ATAC-seq data were processed to call narrow peaks using the nf-core atacseq pipeline v2.1.2 [79,80] with default parameters and read lengths corresponding to the FASTQ files specified at 25, 50 and 75 bp for dog, human and mouse, respectively. This pipeline, built in Nextflow v24.04.2, follows the ENCODE project ATAC-seq processing standards [127]. Raw read files were processed through the pipeline comprising quality control with FastQC v0.11.9 [128] and Cutadapt v3.4 [129], sequence alignment to the reference genome using BWA-MEM v0.7.17 [130], and duplicate marking with Picard v3.0.0 [131]. Alignments from multiple libraries or replicates were merged where required. Alignments were then filtered using SAMtools v1.15.1 (filtering) and v1.17 (sorting, indexing and statistics) [132], BEDTools v2.30.0 [133] and BamTools v2.5.2 [134] to remove duplicate, unmapped, low-quality, and mitochondrial reads. Alignment quality was assessed using Picard v3.0.0 [131] and depth-normalized BigWig coverage tracks scaled to 1 million mapped reads were generated using BEDTools v2.30.0 [133] and UCSC BedGraphToBigWig v445 [135]. MACS2 v2.2.7.1 [136] was used to call narrow peaks, which were annotated relative to gene features using HOMER v4.11 [137]. BEDTools v2.30.0 [133] was then used to create consensus peaks across all samples. Reads in consensus peaks were counted via featureCounts v2.0.1 [138] and differential accessibility analysis, principal component analysis (PCA), and clustering were conducted in R v4.0.3 using DESeq2 v1.28.0 [139]. Finally, results were presented in the form of an IGV session [140] and the whole pipeline was quality checked using ataqv v1.3.1 [141] and MultiQC [142]. For each of the three species, called peaks from all three tissues were then combined into one dataset.

### ExplaiNN: testing on mouse

The mouse was selected as the focal species for the initial ExplaiNN models because it had an intermediate number of samples (18, compared with 29 for human and 10 for dog). Each ExplaiNN run required binary labelled sequences based on chromatin accessibility. We defined ‘positive’ sequences as those intersecting with ATAC-seq peaks, representing putative functional regulatory elements that provide the feature patterns for model training. Conversely, ‘negative’ sequences were sampled from non-coding regions lacking detectable ATAC-seq signals across the three surveyed tissues (heart, stomach, liver). Although latent regulatory activity in other tissues cannot be ruled out, these sequences serve as background non-coding controls which do not have a known function.

Following the ExplaiNN documentation (https://github.com/wassermanlab/ExplaiNN), mouse peak summits from chromosomes 2-19 were extended by ± 100 bp, resulting in 201 bp ‘positive’ sequences. Chromosome 1 was removed at this stage to enable it to be used as an independent test set. The 201 bp ‘positive’ sequences were subsampled to 10,000 sequences using subsample-seqs-by-gc.py [143]. An equivalent number of ‘negative’ sequences were generated by dinucleotide shuffling of the ‘positive’ sequences using BiasAway [144]. The data were split into training, validation and test sets (80/10/10%, respectively) using fasta2explainn.py [143]. ExplaiNN uses the three sets as follows: the model is fitted on the training set, performance on the validation set is monitored during training so that training stops once validation performance no longer improves, and the final model is evaluated on the test set. An ExplaiNN model (Model 1; Table 1) was trained using train.py [143] employing binary cross-entropy with logits (BCEWithLogitsLoss) as the loss criterion and incorporating reverse complement sequences during training. The pipeline output the trained model, training and validation loss metrics, and a parameter file. The trained model and parameter file were provided as input to test.py [143] along with the test dataset (10% of the original dataset) and performance metrics were calculated and provided as output. Model performance was evaluated using the area under the receiver operating characteristic curve (AUC-ROC) and the area under the precision-recall curve (AUC-PR).

A new model (Model 2; Table 1) was trained to ensure the model learned to distinguish between sequences that overlap ATAC-seq peaks and those that do not, rather than real DNA and dinucleotide-shuffled DNA. ‘Positive’ sequences from the prior run were used to train Model 2. ‘Negative’ sequences for Model 2 were generated by defining 201 bp windows across the genome, excluding windows that overlapped with annotated features or ATAC-seq peaks using BEDTools v2.31.1 intersect [133] and then hard-masking the remaining windows by converting soft-masked repetitive sequences to Ns. Any window containing ≥ 1 N was then removed, leaving only sequences with no repetitive regions. As there were far more ‘negative’ than ‘positive’ sequences, a set of ‘negative’ sequences was randomly extracted using “shuf” to approximately match the number of ‘positive’ sequences. Additionally, ‘positive’ and ‘negative’ sequences from chromosome 1 were separated from the main dataset to form an additional test set. The ‘positive’ sequences and a subset of the ‘negative’ sequences from the remaining chromosomes were then analysed with match-seqs-by-gc.py [143] to extract 100,000 ‘positive’ and ‘negative’ sequences matched by GC content. From these, training/test/validation datasets were generated (80/10/10%) and Model 2 was trained. Model 2 was then tested on the set of ‘positive’ and ‘negative’ sequences from chromosome 1.

Equivalent models were trained on the human (Model 3; Table 1) and domestic dog (Model 4; Table 1), as well as a combined human and mouse model (Model 5; Table 1). Again, for each species, chromosome 1 was arbitrarily selected and separated from the rest of the genome to act as an independent test set and the remaining chromosomes were used to generate a training, test, and validation dataset. Training, test, and validation sets were generated using 201 bp peak-summit-flanking sequences as ‘positives’ and 201 bp unannotated non-ATAC-peak genomic windows as ‘negatives’. The same was done for chromosome 1 to generate a labelled independent test set. Once trained, Models 3 and 4 were run on their corresponding chromosome 1 test set. For the combined human and mouse model (Model 5), the existing test sets, generated as described above, were concatenated to ensure equal representation from each species.

### ExplaiNN: transferring between species

After calculating model performance in scenarios where both training and testing data were derived from the same species, models trained on one species were then tested on another species. Each single-species model (mouse, Model 2; human, Model 3; dog, Model 4; Table 1) was tested against the chromosome 1 test set of the other two species and model performance was computed. The human and mouse combined model (Model 5) was tested on the human and mouse combined chromosome 1 test set and the dog chromosome 1 test set.

ExplaiNN Models 1-5 initially made predictions on 201 bp sequences which were either ‘positive’ ATAC-seq peaks or ‘negative’ non-coding, non-repetitive, non-ATAC-seq windows. To test model performance in a non-model species genome, where ‘positives’ and ‘negatives’ are unknown, it was necessary to investigate model performance when predicting on unlabelled genomic windows.

A new chromosome 1 test dataset for each model species was generated by calling 201 bp windows with 151 bp overlap across the whole chromosome using SeqKit v0.10.0 [145]. Genomic windows were labelled as ‘positives’ if they shared at least 75% reciprocal overlap with one of the previously defined 201 bp ATAC-seq peak-summit-flanking positive sequences, to ensure each sequence was assigned as a positive only once. ‘Negative’ windows were defined as non-coding regions showing no overlap with ATAC-seq peaks, annotated coding exons, or repetitive elements. Models 2, 3 and 4 were subsequently run on these positive and negative windows. To assess model performance, AUC-PR and AUC-ROC were calculated. Due to the severe imbalance in the test datasets (far more ‘negatives’ than ‘positives’), AUC-PR and AUC-ROC may not be the most informative metrics [146,147]. Therefore, precision, recall and F1 scores were also calculated [146] and the number of true and false positives and negatives at different classification thresholds was quantified in the dog, as it had the least available data and thus acted as a ‘test’ non-model species.

### ExplaiNN: reducing number of false positives

Methods of differentiating between true- and false-positive predictions were explored with the aim of increasing confidence in the predicted functional regions in the cheetah. Using per-base PhyloP conservation scores in the domestic dog genome [148], the relationship between sequence conservation and ATAC-seq peaks was investigated. Using liftOver (KentUtils v1.0) [135] and the corresponding UCSC chain file, PhyloP scores were converted from canFam4 (GCA_011100685.1) to ROS_Cfam_1.0 (GCA_014441545.1). Predictions from the dog-trained and -tested model (Model 4) were classified based on both the ExplaiNN prediction score and positive/negative labels. The four classification categories were: true positives (TP; labelled as positives, ExplaiNN prediction ≥ 0.5), false positives (FP; labelled as negatives, ExplaiNN prediction ≥ 0.5), true negatives (TN; labelled as negatives, ExplaiNN prediction < 0.5) and false negatives (FN; labelled as positives, ExplaiNN prediction < 0.5). The median PhyloP score for each 201 bp window was calculated and medians plotted for each of the four classification categories. Additionally, the distance from the end of each window to the nearest downstream coding sequence (CDS) was calculated using BEDTools v2.31.1 closest [133] and plotted for each of the four classification categories. Pairwise Mann–Whitney U tests were run on each pair of categories using SciPy Stats [149].

We further evaluated the convolutional filters learned by ExplaiNN to identify those contributing most to model predictions. A filter refers to a 19-bp DNA motif generated by ExplaiNN as a predictive feature. For each of the three species, interpret.py [143] was deployed on the chromosome 2+ model (Models 2, 3 and 4) using the corresponding test dataset (a subset of the chromosome 2+ set), which outputs position weight matrices (PWMs) representing the top 100 19-bp filters in MEME format. We then used meme2logo.py to turn each PWM into a sequence logo. Transcription factor (TF) motifs, represented as position frequency matrices, were downloaded in MEME format from the JASPAR 2024 CORE vertebrates non-redundant collection (https://jaspar.elixir.no/downloads/). Tomtom.py was then used to identify any of the ExplaiNN filters that significantly matched (q-value < 0.05) a JASPAR motif. For each model, the predictive importance of each filter that matched a TF motif was extracted from ExplaiNN interpret.py output and a heatmap of these values was plotted using the Python package seaborn [150].

### ExplaiNN: running on cheetah and comparing to sequence conservation

A final ExplaiNN model (Model 6) was trained on a total of 100,000 positive and 100,000 negative sequences taken from the human, mouse and domestic dog datasets, with approximately equal representation from each species. The cheetah genome (VMU_Ajub_asm_v1.0, GCA_027475565.2, [94]) was split into 201 bp sliding windows with a 151 bp overlap. The trained Model 6 was then deployed on these cheetah windows to predict functional non-coding regions.

To compare model predictions with sequence conservation, PhyloP scores were calculated and ultra-conserved elements (UCEs) extracted. A genome alignment comprising thirteen felids and four mammalian outgroups (Table S3) was generated using Cactus v2.9.8-gpu [151]. Genomes were selected based on chromosome-level assembly, availability of a RefSeq annotation, and to ensure even taxonomic sampling across Felidae. From the HAL file generated by Cactus, the ancestral genome (Anc00) was extracted using HAL v2.2 hal2fasta [152]. Using SeqKit v2.1 [145], the ancestral genome was unmasked and re-masked using RepeatMasker v4.0.8 [153] to identify ancestral repeats. Ancestral repeats were extracted from the cheetah genome using HAL v2.2 Liftover [152] and filtered to remove any complex repeat regions or coding regions. A MAF file was generated using Cactus v2.9.8-gpu hal2maf [151] with the cheetah as a reference. From this, a neutral substitution model was generated using phyloFit v1.4 [63] with a species tree extracted from existing literature (Table S3; [154,155]). Using KentUtils v1.0 mafSplit [135], the MAF file was separated into chromosomes. We used BEDTools v2.31.1 makewindows [133] to bin the cheetah genome into 25 bp windows. Using PhyloP v1.4 [63] and the fitted neutral model, we calculated PhyloP scores in 25 bp windows across the 19 chromosomal scaffolds of the cheetah genome under the conservation/acceleration mode (--mode CONACC) using the likelihood ratio test (--method LRT). The majority of smaller contigs did not have sufficient alignment to calculate conservation scores, so all non-chromosomal contigs were discarded at this stage. The mean conservation score for each predicted functional non-coding region in the chromosomes was calculated using the PhyloP scores. The neutral model generated by phyloFit was also provided to PhastCons v1.6 [156] with the “--most-conserved” flag to identify ultra-conserved elements (UCEs).

Based on testing in the dog, we ran several filtering steps on the cheetah genomic windows predicted as positives by ExplaiNN Model 6. Firstly, windows with any overlap to genes were filtered out using BEDTools v2.31.1 intersect [133] to retain only non-coding windows. Next, windows with an ExplaiNN prediction score ≥ 0.75 were extracted to retain only those windows with confident positive predictions. The distance from each window to the nearest downstream gene start site was calculated using BEDTools and windows were filtered to retain only those within 50,000 bp of a gene start site, as these regions are more likely to contain promoters or enhancers [88–93]. Finally, remaining windows were filtered based on mean PhyloP scores to retain only windows with a mean score ≥ 2, suggesting significant evolutionary constraint compared to the neutral model [1,157].

To compare the resultant set of predicted functional non-coding regions to alternative methods of annotating the non-coding genome, UCEs were extracted from the MAF file using PhastCons v1.6 [156]. To generate a conservative set of high-confidence functional non-coding regions, we filtered genomic windows using three stringent criteria: an ExplaiNN score ≥ 0.75, proximity to nearest downstream gene (≤ 50 kb), and evidence of evolutionary constraint (PhyloP score or overlap with a UCE). Additionally, FIMO (MEME suite v5.1.0) [158] was used with default settings to scan the resultant set of windows for transcription factor binding sites using the JASPAR 2024 Vertebrate Core Non-redundant motif database (https://jaspar.elixir.no/downloads/).

### Variant annotation in the cheetah

We called variants from whole-genome resequencing data generated from 39 cheetahs, comprising 30 individuals from the U.S. *ex situ* population and nine wild-origin individuals from Tanzania (n=3), Namibia (n=4), and South Sudan (n=2). Full details of sample collection, sequencing, and quality control are provided in [69]. SNPs were called against the VMU_Ajub_asm_v1.0 cheetah assembly (GCA_027475565.2, [94]) and filtered using default GATK v4.6.0.0 hard filtering parameters [159–161] with a reduced FS (FisherStrand) requirement (FS < 31.6; 99th percentile) corresponding to our data, and a minimum and maximum per-site depth of 119 (1st percentile) and 837 (99th percentile), respectively (see [69] for full methods). SNPs occurring within the 201 bp putative functional non-coding windows predicted by ExplaiNN were extracted using BEDTools v2.31.1 intersect [133].

Because tools like SnpEff [162] are not applicable to predict the deleteriousness of non-coding variants, several other methods were applied to predict the functional impact of these SNPs. Combined Annotation-Dependent Depletion (CADD) scores quantify the potential deleteriousness of genetic variants using genomic annotations to predict their functional impact. Human CADD scores (hCADD) were downloaded from CADD v1.7 [62] and lifted over from the human to the cheetah reference genome using LoadLift [163]. LoadLift only transfers scores within ultra-conserved elements (UCEs), where sequence conservation makes the effects of mutations more likely to be similar between species. As a result, SNPs in predicted functional non-coding windows outside UCEs did not receive a CADD score. To increase coverage, precomputed NCBoost scores [164] were also applied, which use supervised learning with gradient tree boosting to assess pathogenic non-coding variants associated with monogenic Mendelian diseases. To map these scores onto the cheetah genome, a human-to-cheetah chain file was generated using Cactus v2.9.8-gpu hal2chains on the previously generated 17-species HAL alignment file [151,152]. Using this, pre-computed NCBoost scores for >850 million human genomic positions were lifted over to the cheetah genome with UCSC LiftOver [135,164]. CADD and NCBoost scores overlapping each SNP were extracted using BEDTools v2.31.1 intersect [133]. SNPs with CADD or NCBoost scores above the 75th percentile of their respective score distributions were retained. These SNPs were intersected with significant FIMO (MEME suite v5.1.0) [158] hits using BEDTools v2.31.1 intersect [133]. SNPs meeting both criteria were designated as putatively deleterious non-coding SNPs.

To test for associations between this SNP set and genes they may impact, cheetah Hi-C data were used to define topologically associating domains (TADs). A whole-blood sample from a male cheetah (studbook number AJU9890) housed at Toronto Zoo was collected during a routine veterinary exam in EDTA Vacutainer® tubes (Becton Dickinson, U.S.). Library preparation and Arima Genomics High Coverage Hi-C sequencing were completed by MedGenome. We processed raw Hi-C reads using Juicer v1.6 [165], aligning them to the VMU_Ajub_asm_v1.0 assembly (GCA_027475565.2; [94]) with BWA v0.7.17 [130] under the Arima Genomics Hi-C restriction site configuration. PCR and optical duplicates were removed, and reads were filtered to retain only alignments with MAPQ ≥ 30. TADs were called using Juicer v1.6 Arrowhead [165] with resolutions of 10, 25 and 50 kb, with the latter used for subsequent analysis.

For each SNP, the nearest downstream gene within the same 50 kb-resolution TAD was identified using BEDTools v2.31.0 closest [133], forming a ‘TAD-informed’ gene set. For SNPs with no downstream gene within the TAD, the nearest upstream gene was taken instead. As not every SNP fell within a TAD, the nearest downstream gene for each SNP was identified using BEDTools closest in a strand-aware manner, forming a ‘proximity-informed’ gene set.

For both gene sets, gene names were converted to the Ensembl gene IDs of their human orthologues using Ensembl BioMart v115 [166]. Those with a corresponding 1:1 ortholog in the human were retained and used as the foreground for Gene Ontology (GO) enrichment analysis. We ran OrthoFinder v2.5.4 [167] on the longest protein isoform of each gene in the most recent cheetah, human, domestic dog and domestic cat reference assemblies (GCA_027475565.2, GCA_000001405.29, GCA_014441545.1, GCA_018350175.1, respectively). Genes with a one-to-one (1:1) orthologue between human and cheetah were used as the background gene set. GO enrichment analysis was run for each set of genes (‘TAD-informed’ and ‘proximity-informed’) separately and in combination using ShinyGO v0.86 [168] with an enrichment false discovery rate (FDR)-corrected q-value cutoff of 0.05.

Genes with previously reported deleterious mutations in the cheetah were extracted from the literature [66,69,77,78,119], allowing us to identify genes potentially affected by both coding and non-coding deleterious variation (Table S4). The ‘TAD-informed’ and ‘proximity-informed’ gene lists were combined and manually compared to the literature-derived genes to determine the number of studies reporting each gene. Gene IDs were extracted as described above using Ensembl BioMart v115 [166] and GO enrichment was run on each set of genes using ShinyGO v0.86 [168] as described above.

Unique, putatively deleterious non-coding SNPs were extracted for each population to determine the proportion of unique non-coding mutations. For each sample, we calculated the proportion of putatively deleterious non-coding SNPs with a called genotype that were homozygous for the alternate allele or heterozygous. Kruskal–Wallis tests were carried out using SciPy Stats [149] to assess whether these proportions differed significantly across *ex situ* and wild populations and pairwise Mann–Whitney U tests with Bonferroni correction were carried out using SciPy Stats [149] to identify which pairs of populations differed significantly. These proportions were visualised with Matplotlib [169].

## Declarations

### Ethics approval and consent to participate

The sample for the cheetah Hi-C data was collected opportunistically (e.g., during routine veterinary practices) and provided with full permission from Toronto Zoo.

### Consent for publication

Not applicable.

### Availability of data and materials

The majority of data analysed during the current study are publicly available and can be found using the corresponding accession numbers, detailed in the methods, with the exception of the cheetah Hi-C data, which is available in the ENA (PRJEB130886). The CanFam4-referenced per-base PhyloP score files are available on Zenodo at DOIs 10.5281/zenodo.20137136 (https://zenodo.org/records/20137136).

### Competing interests

KPK is currently employed by Colossal Biosciences; however, this work was completed prior to that employment. The authors declare no competing interests.

### Funding

JAP was supported by the UK Research and Innovation Biotechnology and Biological Sciences Research Council (BBSRC) Norwich Research Park Biosciences Doctoral Training Partnership (BB/T008717/1). WH was supported by BBSRC grants BB/X011070/1 (BBS/E/ER/230001A, BBS/E/ER/230001B, BBS/E/ER/230001C), BB/X011089/1 (BBS/E/ER/230002A, BBS/E/ER/230002B), and EPSRC EP/X035913/1. SAS was supported by BBSRC grants BB/X011089/1 (BBS/E/ER/230002B), BB/X011070/1 (BBS/E/ER/230001C) and BB/CCG2220/1. The authors also acknowledge support from the BBSRC Core Capability Grant BB/CCG1720/1. KLT is a Distinguished Professor for the Swedish Research Council (2023-00391), a KAW scholar (2023.0307), and a Torsten Söderberg Academy professor in Medicine 2024.

### Authors’ contributions

WH conceptualised and acquired funding for the project. JAP, WJN and WH developed the methodology. JAP carried out data curation, formal analysis and visualisation, and wrote the first draft of the manuscript. SAS conducted the LoadLift analyses and wrote the corresponding methods. WH and WJN supervised the project. HRS contributed to visualisation. MD and KLT generated the canine PhyloP scores. KPK provided the cheetah sample for Hi-C, and EEA generated the Hi-C sequence data and provided guidance for analysis. All authors edited and approved the final manuscript.

## Acknowledgements

We thank the keepers, veterinary team, and staff at the Toronto Zoo for granting permission and providing the sample used the generate the cheetah Hi-C data.

