## Supplementary Material for "Machine learning applications to predict functional non-coding regions in non-model species"

Supplementary Methods

### Processing ATAC-seq data and calling peaks: domestic dog

Input reads per sample ranged from 22,763,816 to 96,521,458, with 18.3 to 64.2% of these marked as duplicates (Figure S1). Per-sequence GC content showed a roughly normal distribution, suggesting a normal random library (Figure S2). All reads passed initial filtering steps. After adapter trimming, 20,624,461 to 70,507,822 reads remained. Raw mapping efficiency was between 74.4 and 91.7%, but after alignment filtering, 1,805,110 to 5,346,126 mapped reads per sample remained. The low number of mapped reads post-filtering is likely due to high duplication rates and the absence of a mitochondrial sequence in the dog reference genome assembly, which likely caused mitochondrial-derived reads to map ambiguously or with low confidence. Between 623 and 5,084 narrow peaks were called per sample (Figure S3). FRiP (Fraction of Reads in Peaks) scores ranged from 0.05 to 0.24. Filtered narrow peaks were subsequently annotated relative to gene features and a periodicity plot was generated by Picard (<https://github.com/broadinstitute/picard>) to show insert sizes (Figure S4).

### Processing ATAC-seq data and calling peaks: human

Input reads ranged from 13,593,438 to 375,684,903, with 10.9 to 73.0% marked as duplicates (Figure S5). Per-sequence GC content showed a roughly normal distribution for about half of the samples, suggesting a normal random library (Figure S6). All reads passed initial filtering steps. After adapter trimming, 13,585,130 to 375,452,793 reads remained. Following alignment, 0.2 to 6.7% of reads remained unmapped. Between 35,572,576 and 730,577,690 reads per sample remained after merging of libraries and replicates and alignment filtering. Between 37,992 and 340,687 narrow peaks per sample were called (Figure S7). FRiP scores ranged from 0.09 to 0.48, peaks were annotated relative to gene features, and a periodicity plot was generated (Figure S8).

### Processing ATAC-seq data and calling peaks: mouse

Input reads ranged from 15,591,507 to 75,813,558, with 14.6 to 50.0% of these marked as duplicates (Figure S9). Per-sequence GC content showed a roughly normal distribution and all reads passed initial filtering steps (Figure S10). After adapter trimming, 15,523,342 to 75,552,933 reads remained. Following alignment, 0.9 to 4.0% of reads remained unmapped. After merging of libraries and replicates for each sample and alignment filtering, 50,713,172 to 173,331,152 mapped reads remained per sample. Between 17,603 and 70,592 narrow peaks were called per sample (Figure S11). FRiP scores ranged from 0.10 to 0.42. Peaks were subsequently annotated relative to gene features and a periodicity plot was generated (Figure S12).

Supplementary Figures

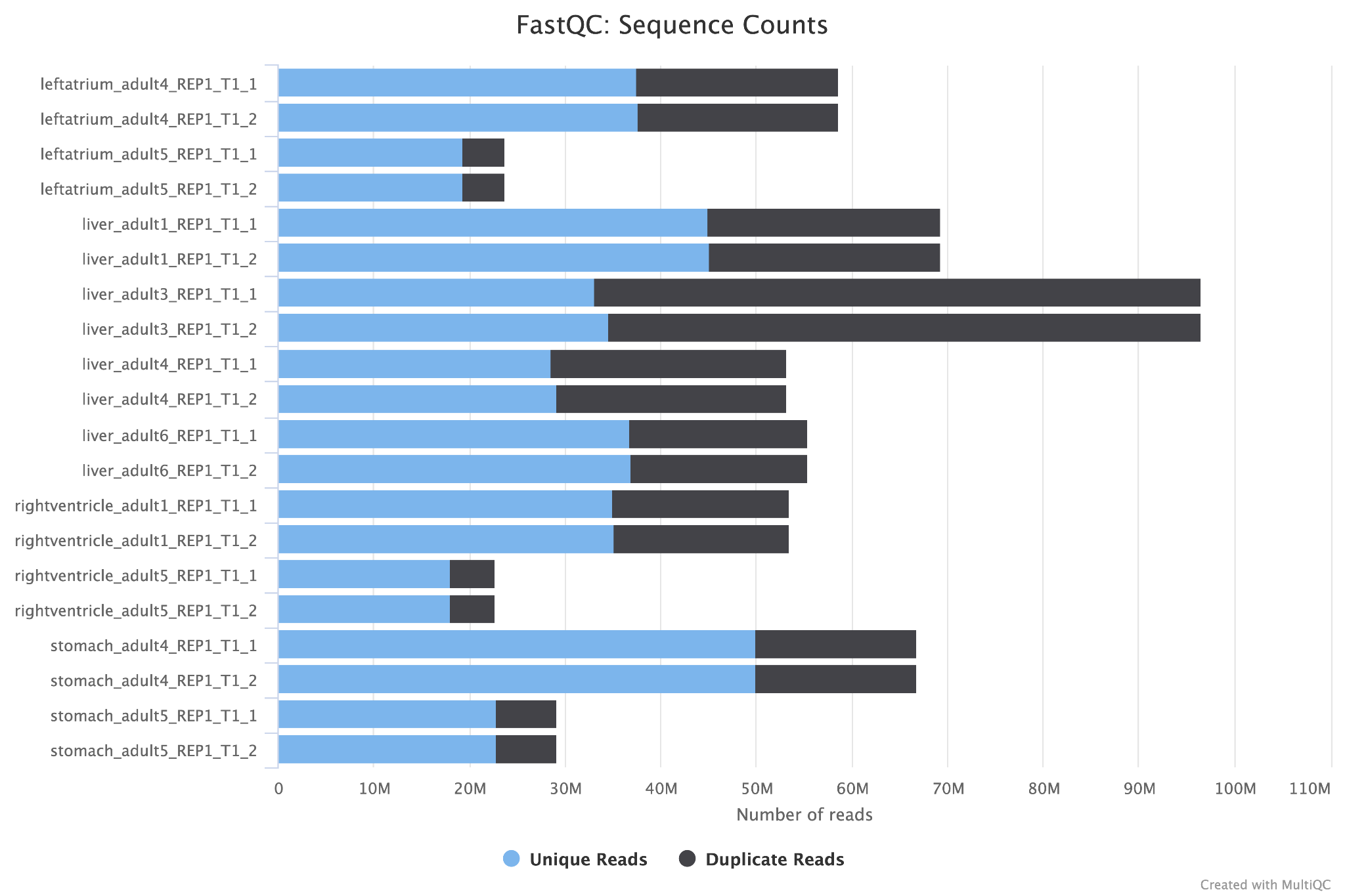

**Figure S1. Total number of ATAC-seq reads in dog.** Total number of reads provided as input for each dog sample (ranging from 22,763,816 to 96,521,458) and proportion of those marked as duplicates (ranging from 18.3 to 64.2%). Plot generated by MultiQC as part of the nf-core atacseq pipeline v2.1.2 (Patel et al. 2023; P. Ewels et al. 2016; P. A. Ewels et al. 2020).

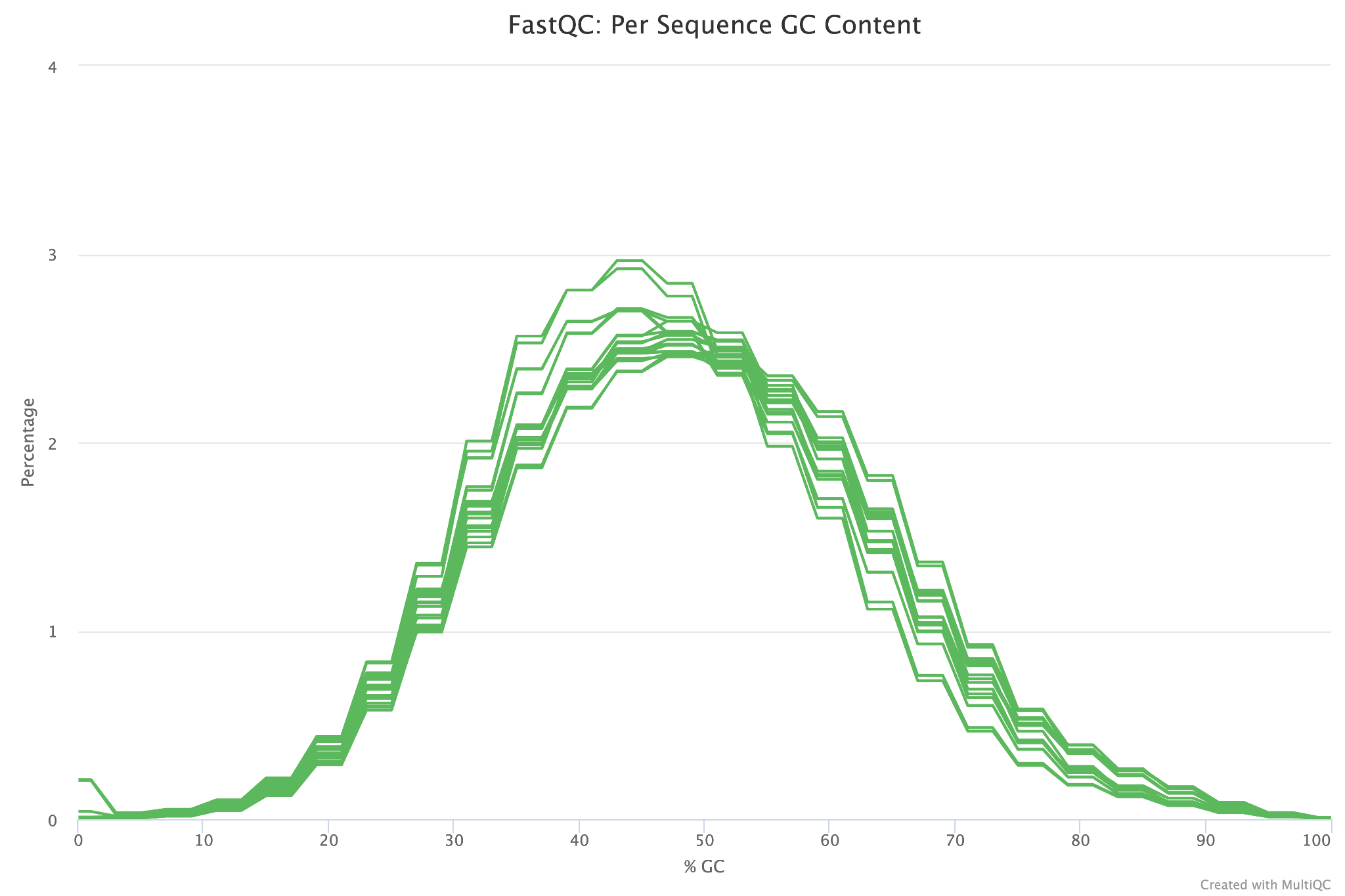

**Figure S2. GC content of ATAC-seq reads in dog.** Per-sequence GC content calculated by FastQC showing roughly normal distribution. Plot generated by MultiQC as part of the nf-core atacseq pipeline v2.1.2 (Patel et al. 2023; P. Ewels et al. 2016; P. A. Ewels et al. 2020).

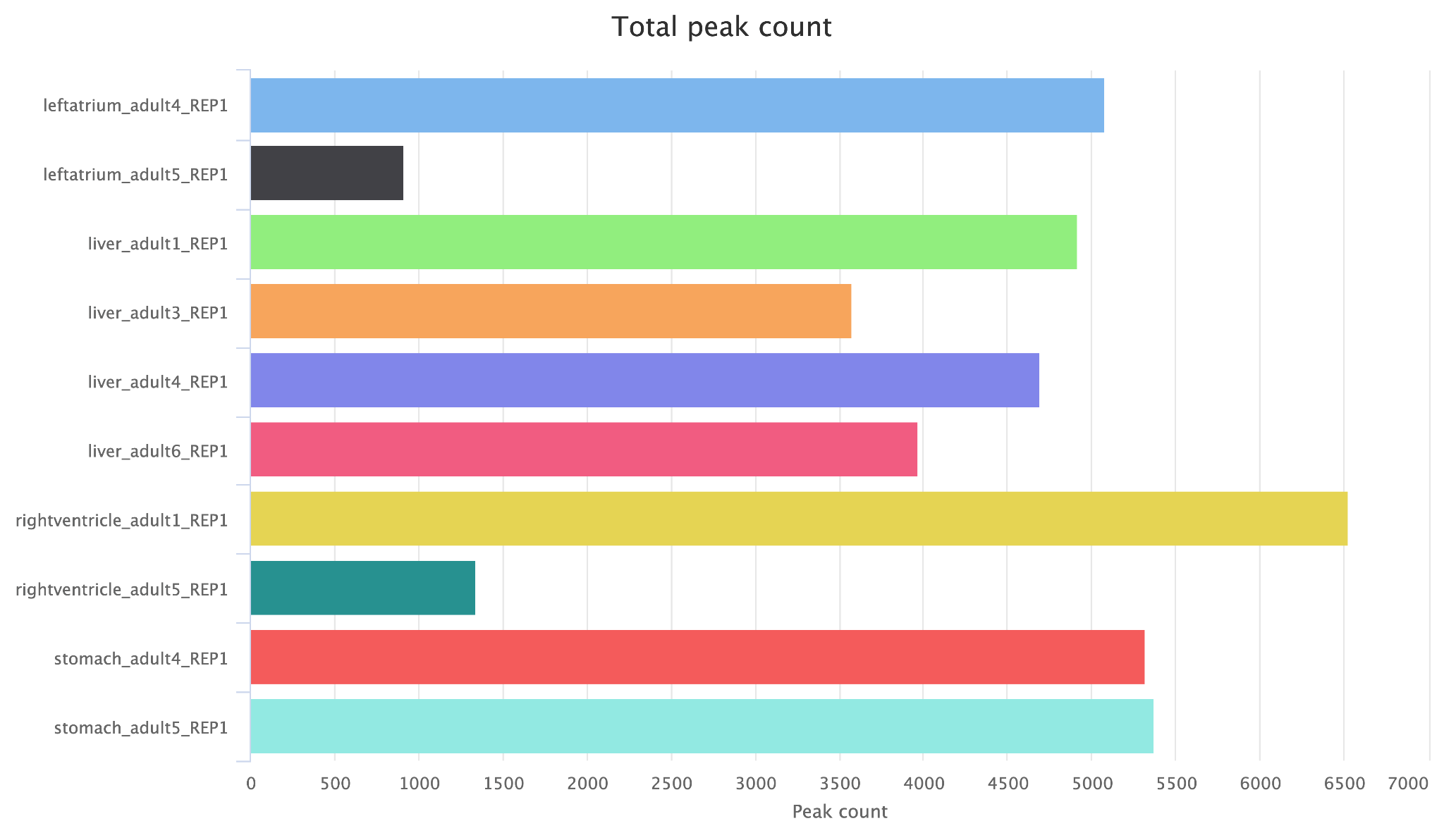

**Figure S3. Total peak count in dog.** Total narrow peak count for each sample as called by MACS2, ranging from 623 to 5,084 peaks per sample. Plot generated by MultiQC as part of the nf-core atacseq pipeline v2.1.2 (Patel et al. 2023; P. Ewels et al. 2016; P. A. Ewels et al. 2020).

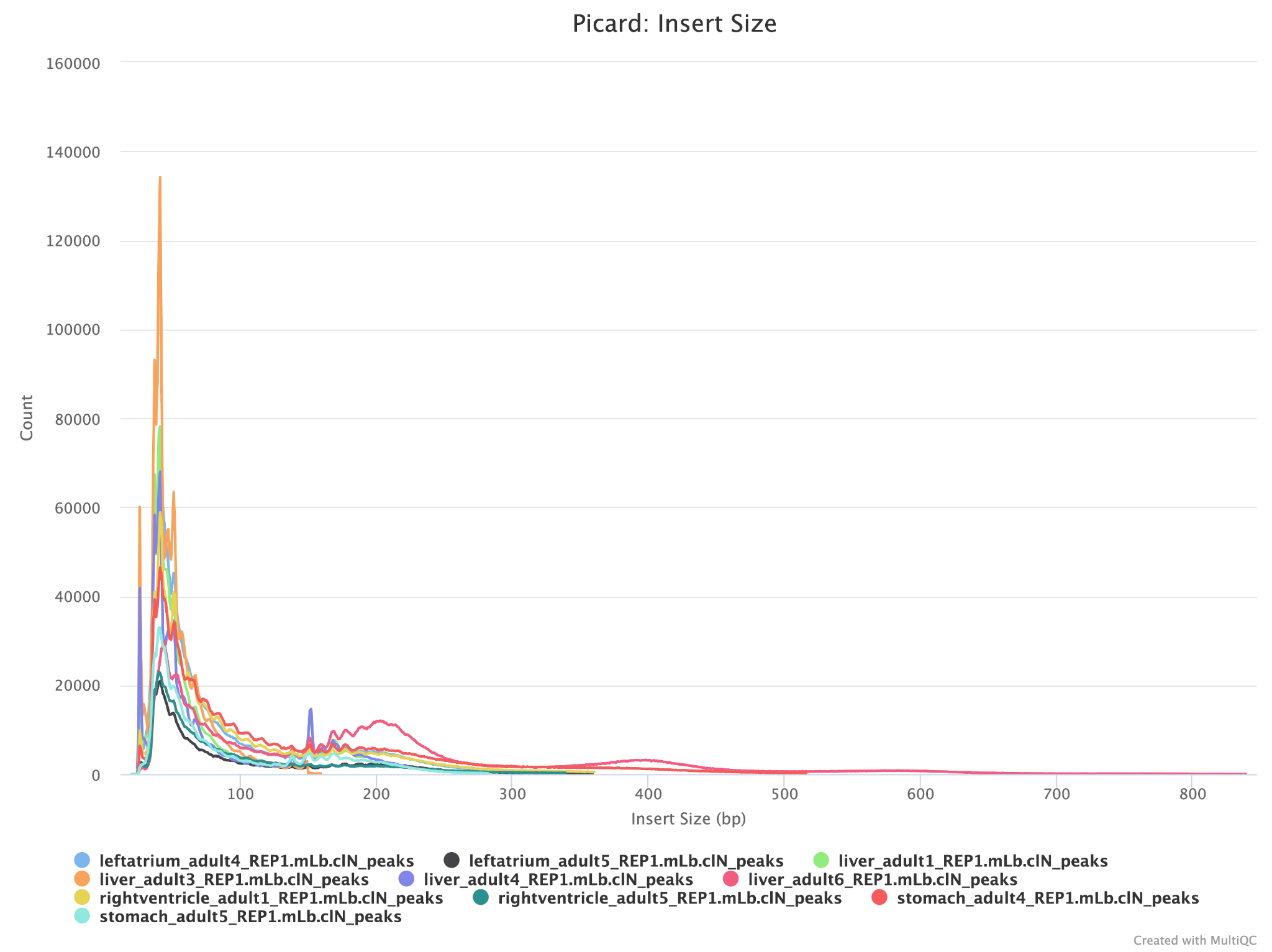

**Figure S4. Periodicity plot in dog.** Periodicity plot showing insert size of each peak. Plot generated by MultiQC as part of the nf-core atacseq pipeline v2.1.2 (Patel et al. 2023; P. Ewels et al. 2016; P. A. Ewels et al. 2020).

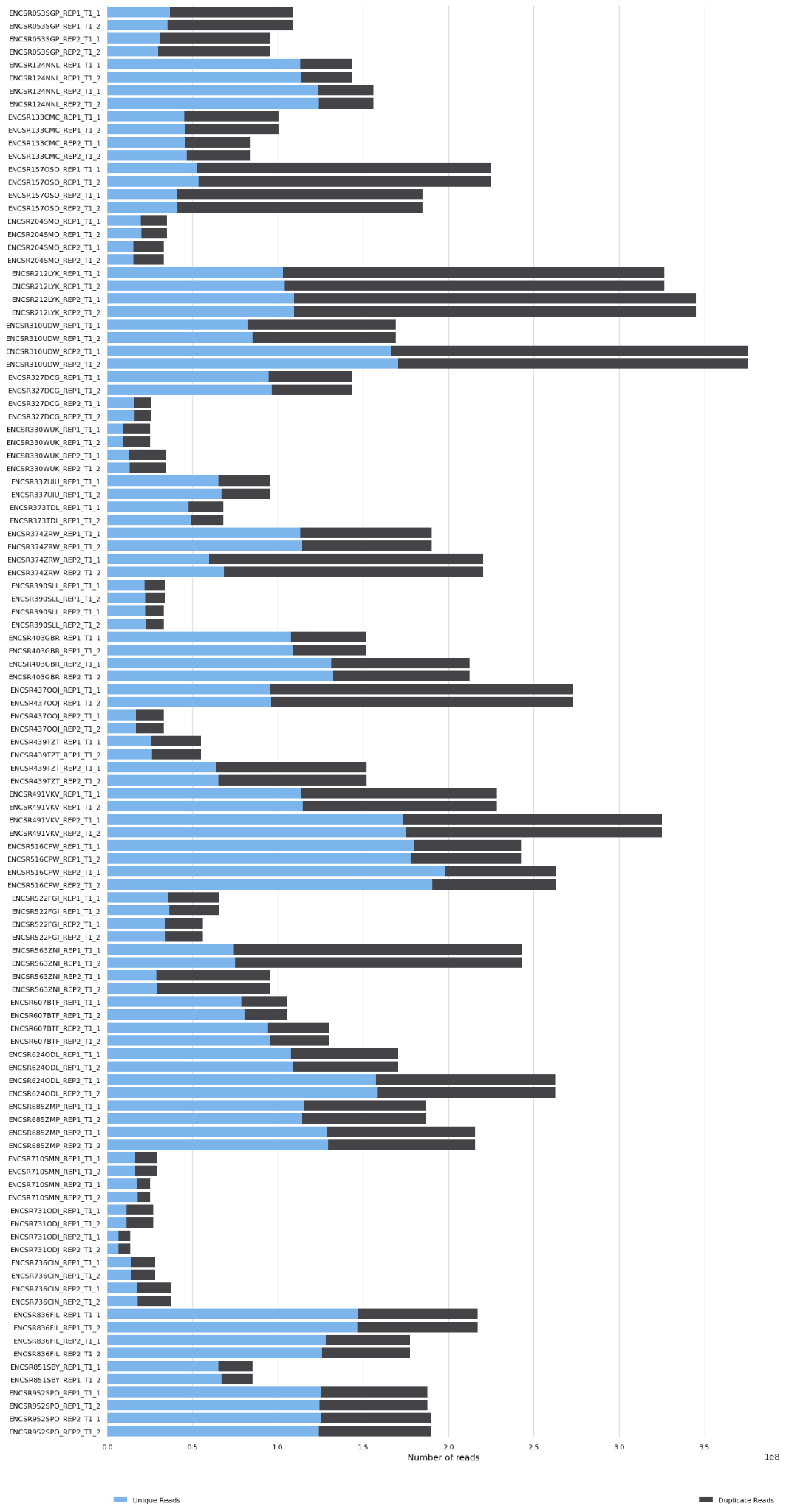

**Figure S5. Total number of ATAC-seq reads in human.** Total number of reads provided as input for each sample (ranging from 13,593,438 to 375,684,903) and proportion of those marked as duplicates. Plot generated by MultiQC as part of the nf-core atacseq pipeline v2.1.2 (Patel et al. 2023; P. Ewels et al. 2016; P. A. Ewels et al. 2020).

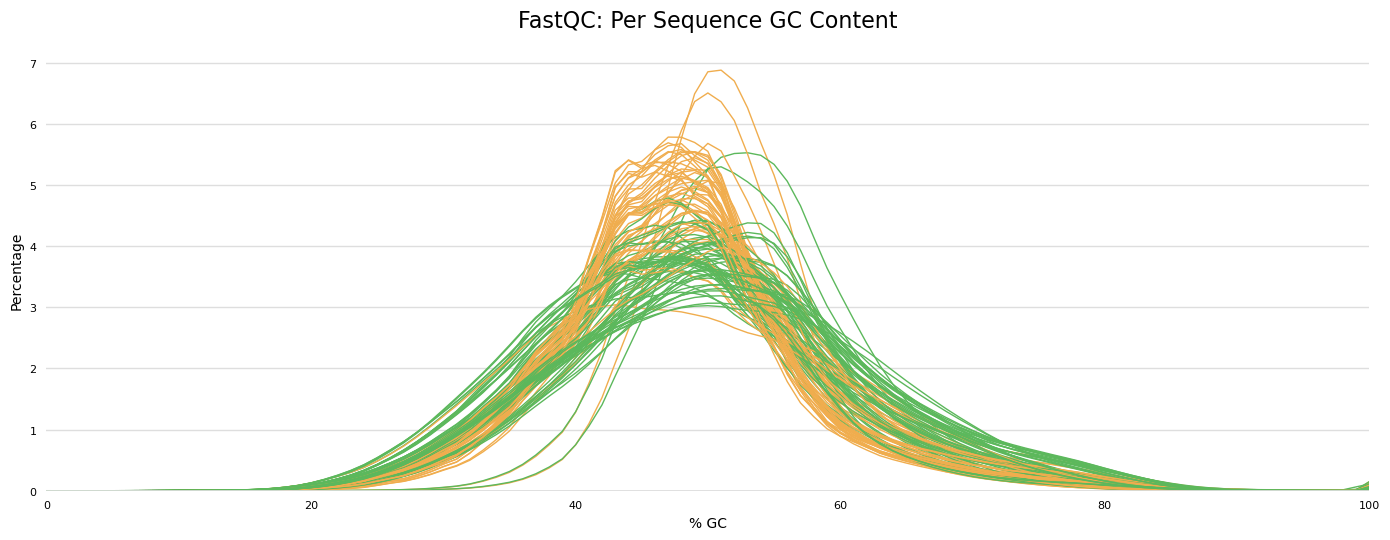

**Figure S6. GC content of ATAC-seq reads in human.** Per-sequence GC content calculated by FastQC showing roughly normal distribution. Plot generated by MultiQC as part of the nf-core atacseq pipeline v2.1.2 (Patel et al. 2023; P. Ewels et al. 2016; P. A. Ewels et al. 2020).

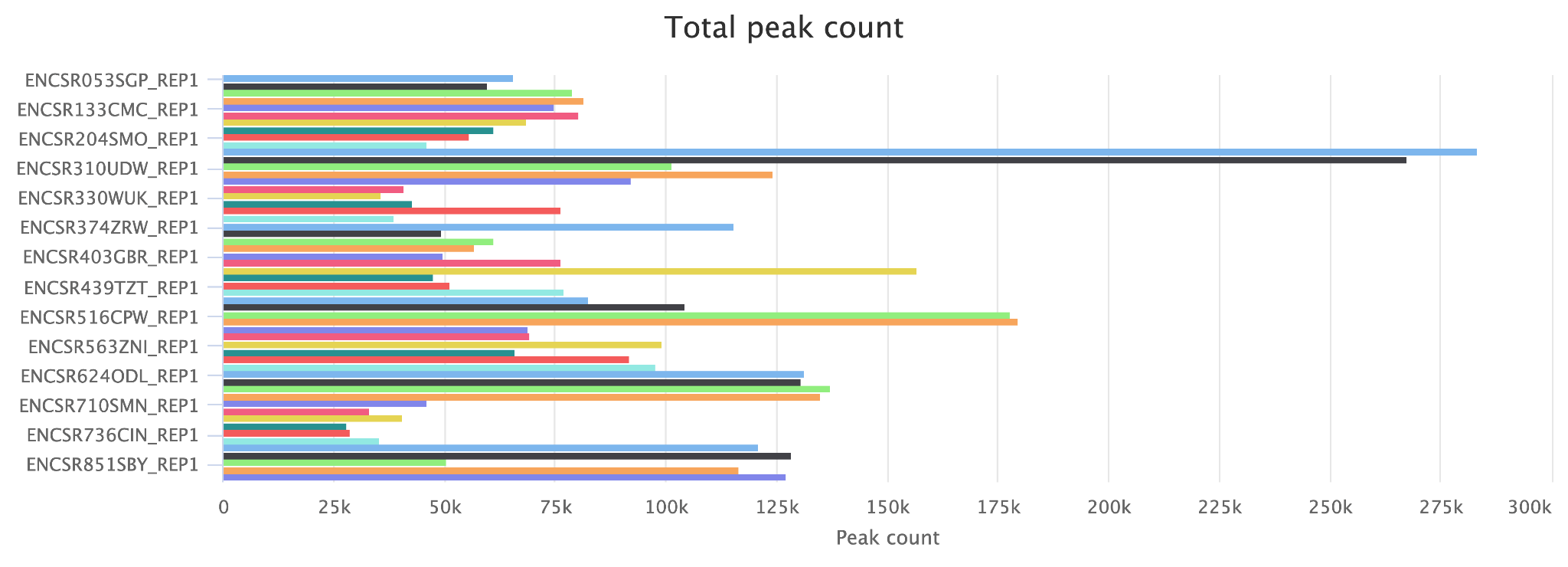

**Figure S7. Total peak count in human.** Total narrow peak count for each sample as called by MACS2, ranging from 37,992 to 340,687 peaks per sample. Plot generated by MultiQC as part of the nf-core atacseq pipeline v2.1.2 (Patel et al. 2023; P. Ewels et al. 2016; P. A. Ewels et al. 2020).

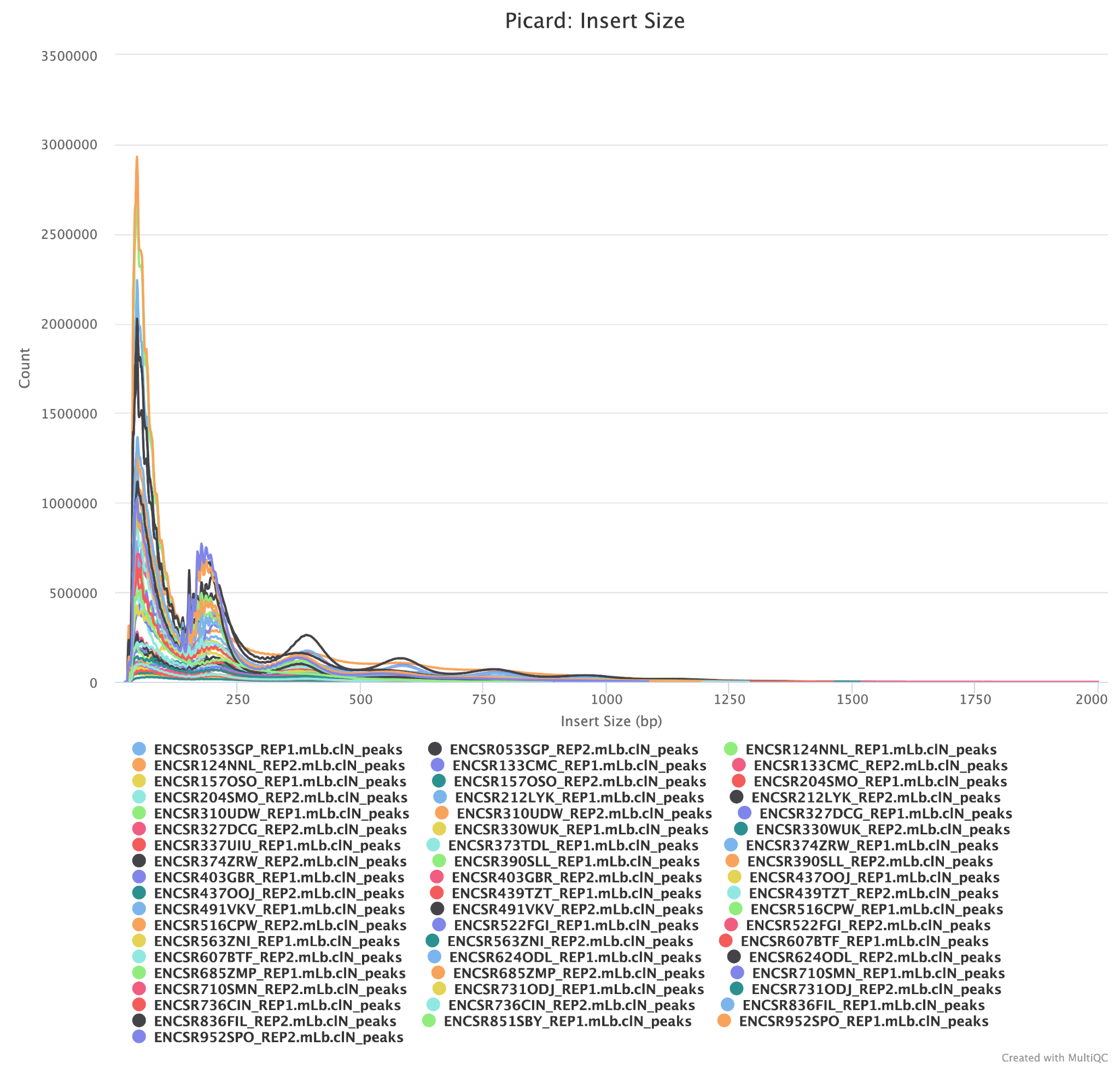

**Figure S8. Periodicity plot in human.** Periodicity plot showing insert size of each peak. Plot generated by MultiQC as part of the nf-core atacseq pipeline v2.1.2 (Patel et al. 2023; P. Ewels et al. 2016; P. A. Ewels et al. 2020).

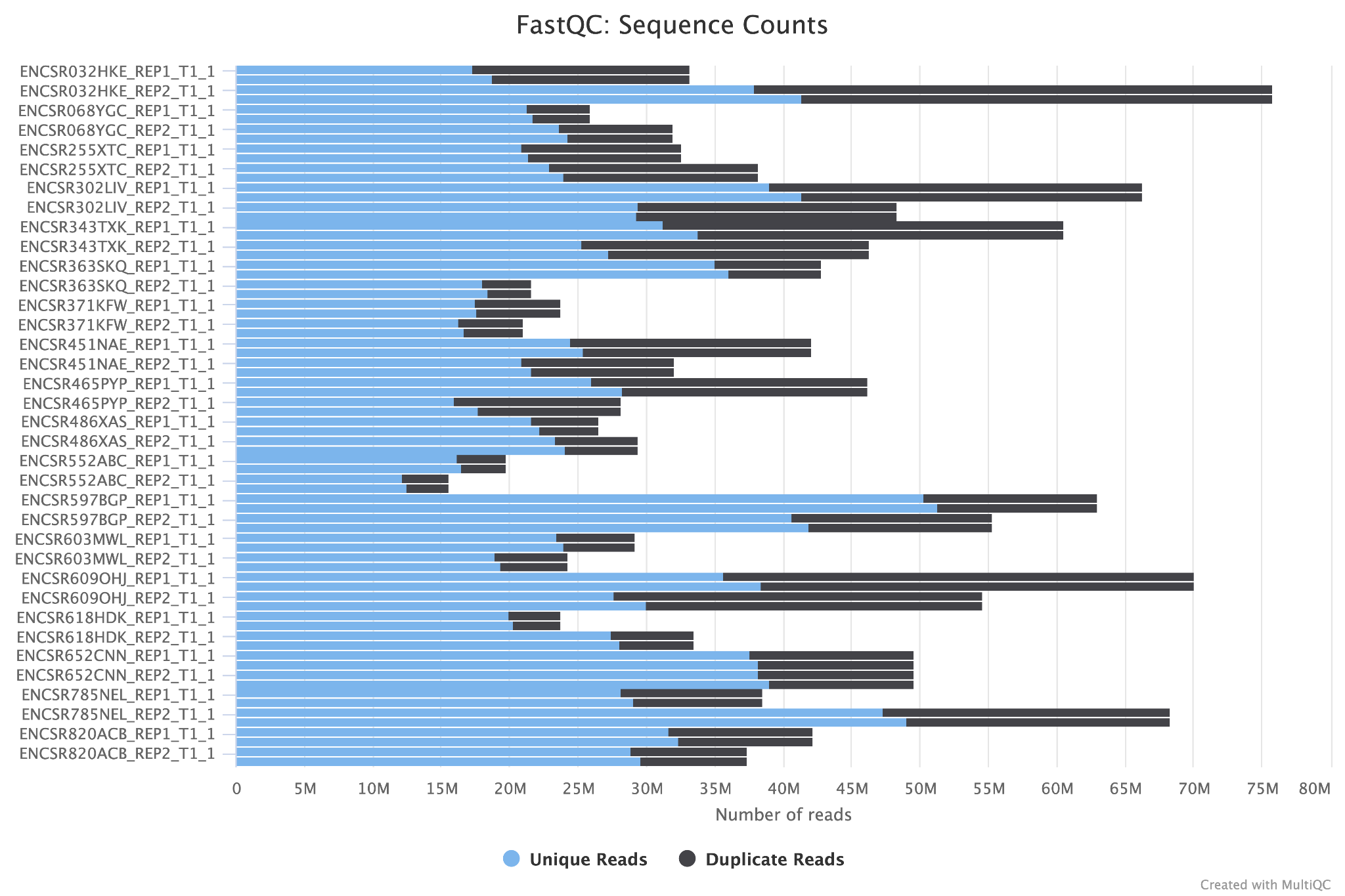

**Figure S9. Total number of ATAC-seq reads in mouse.** Total number of reads provided as input for each sample (ranging from 15,591,507 to 75,813,558) and proportion marked as duplicates (14.6–50.0%). Plot generated by MultiQC as part of the nf-core atacseq pipeline v2.1.2 (Patel et al. 2023; P. Ewels et al. 2016; P. A. Ewels et al. 2020).

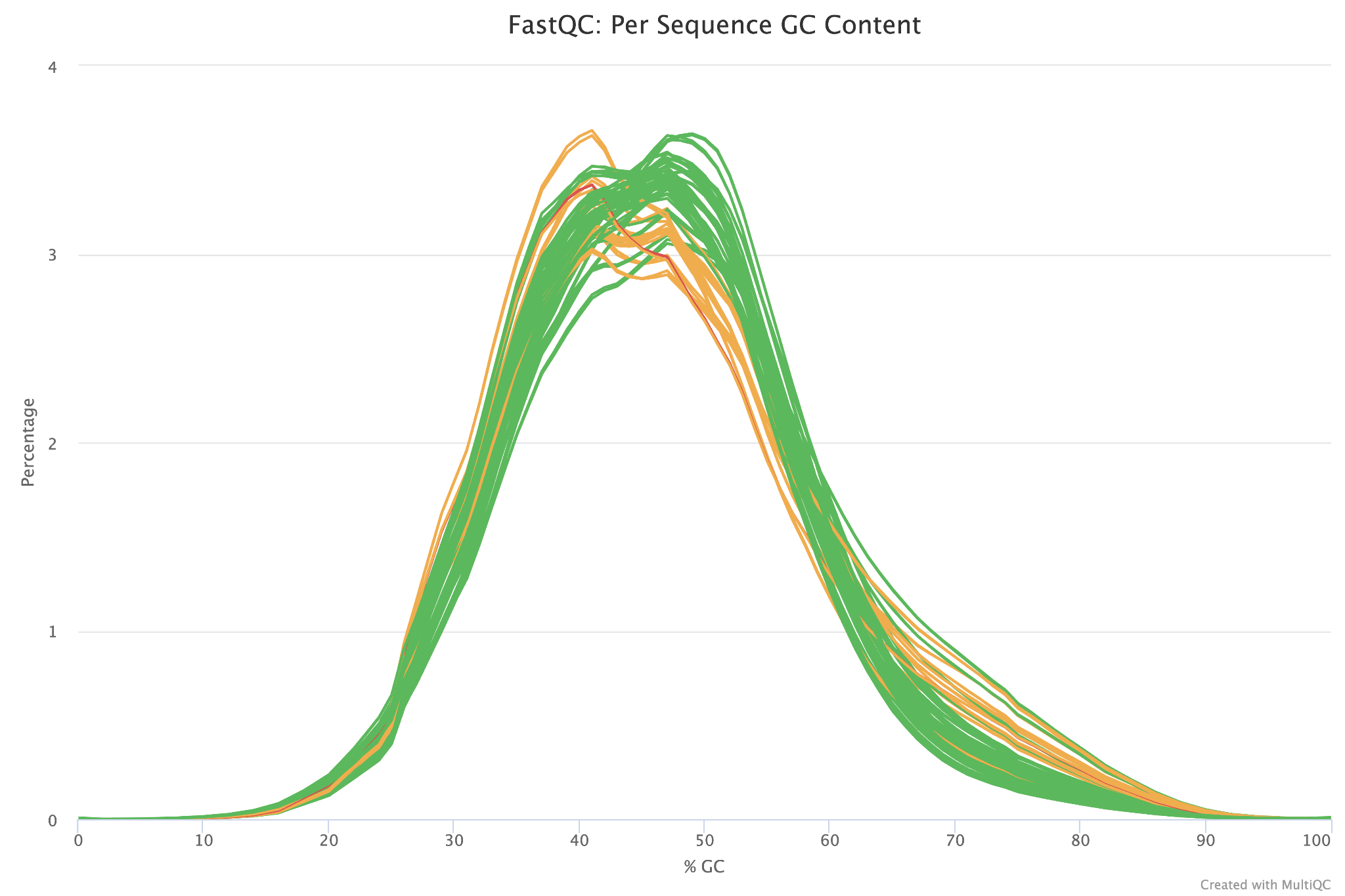

**Figure S10. GC content of ATAC-seq reads in mouse.** Per-sequence GC content calculated by FastQC showing roughly normal distribution. Plot generated by MultiQC as part of the nf-core atacseq pipeline v2.1.2 (Patel et al. 2023; P. Ewels et al. 2016; P. A. Ewels et al. 2020).

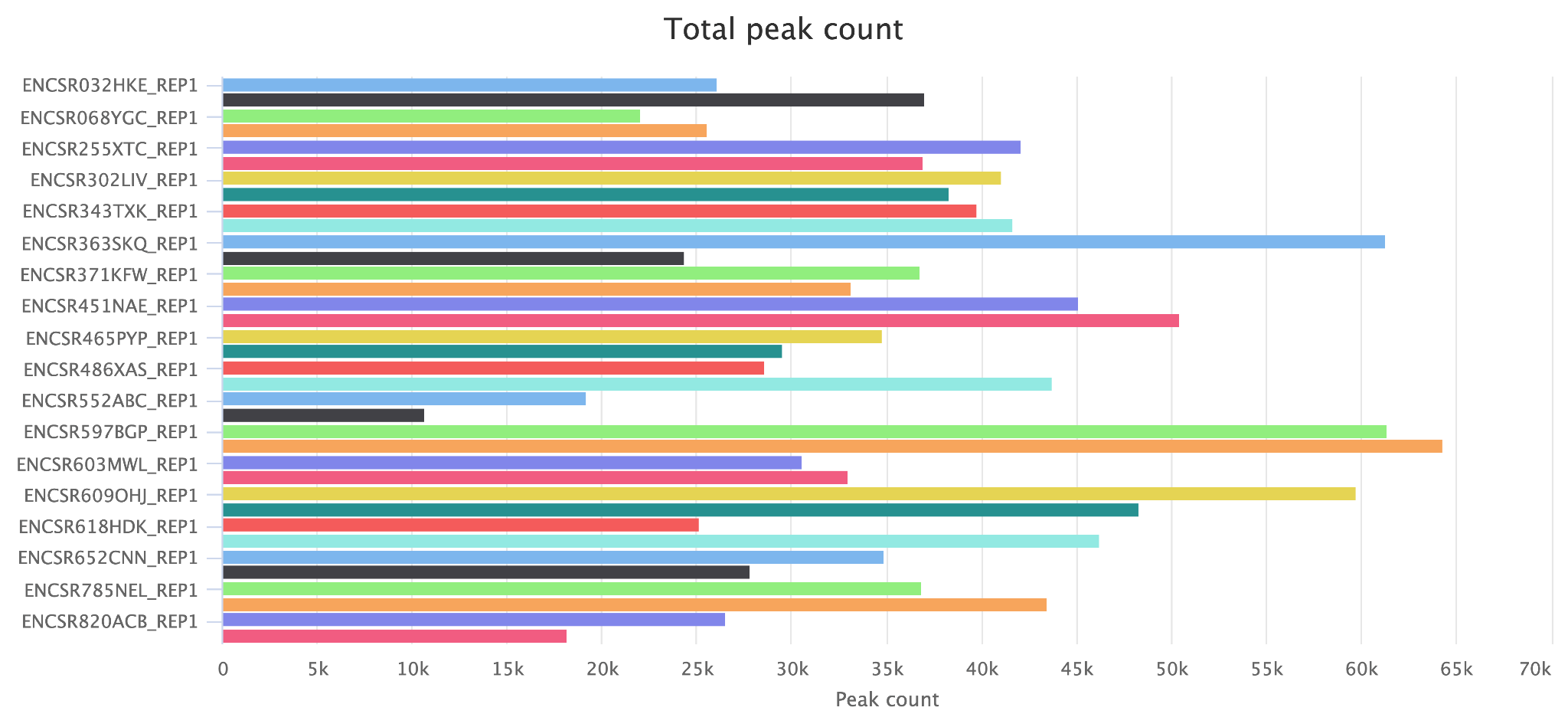

**Figure S11. Total peak count in mouse.** Total narrow peak count for each sample as called by MACS2, ranging from 17,603 to 70,592 peaks per sample. Plot generated by MultiQC as part of the nf-core atacseq pipeline v2.1.2 (Patel et al. 2023; P. Ewels et al. 2016; P. A. Ewels et al. 2020).

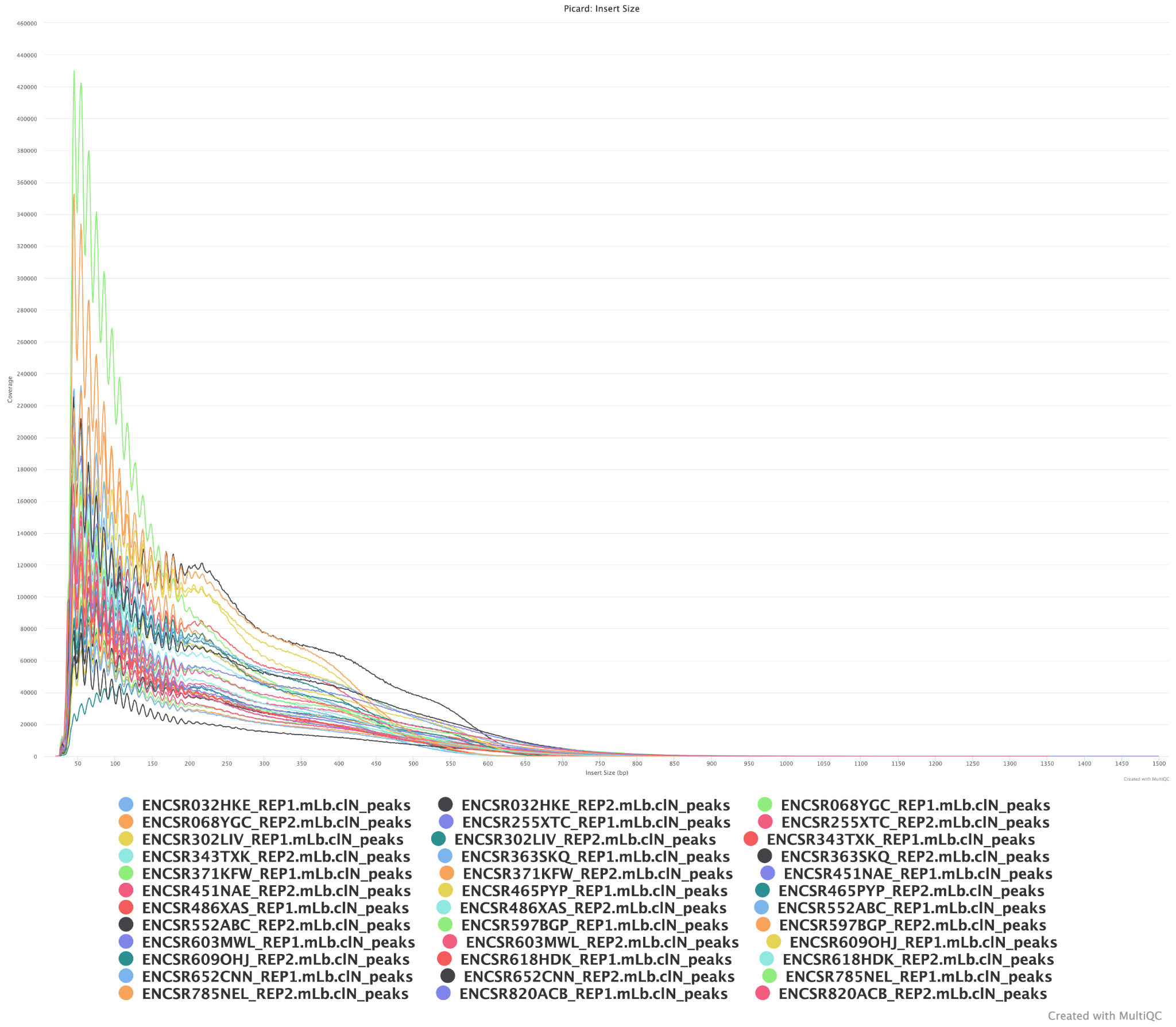

**Figure S12. Periodicity plot in mouse.** Periodicity plot showing insert size of each peak. Plot generated by MultiQC as part of the nf-core atacseq pipeline v2.1.2 (Patel et al. 2023; P. Ewels et al. 2016; P. A. Ewels et al. 2020).

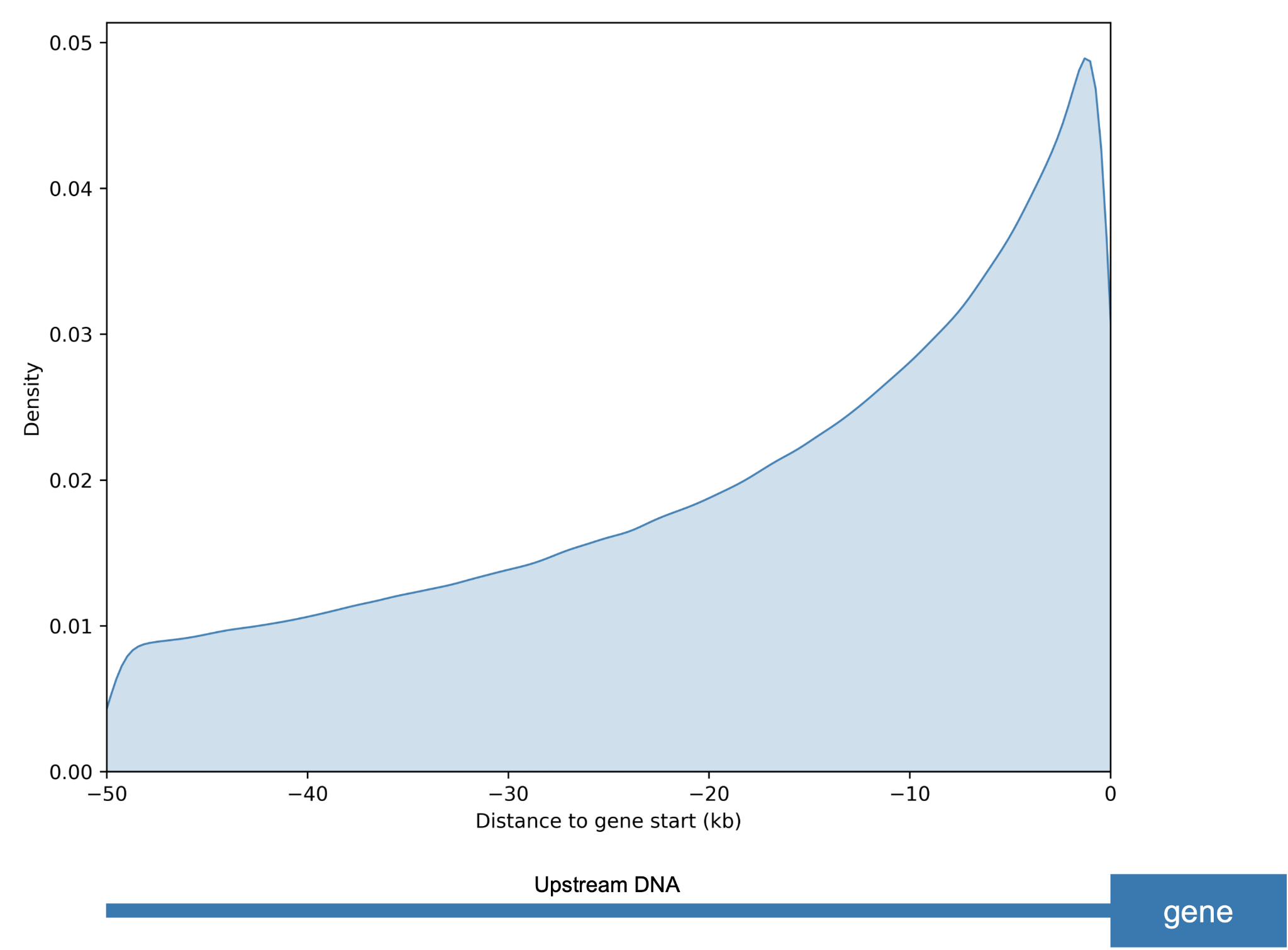

**Figure S13. Distribution of genomic windows relative to gene start sites.** Kernel density estimate (KDE) of distances from 292,912 genomic windows to the nearest gene start site. Distances are shown in kilobases (kb), with negative values corresponding to upstream positions relative to the TSS (0 kb). The density increases toward the gene start site, with the highest concentration of windows located within approximately 5 kb upstream. The schematic below the x-axis illustrates the orientation of the upstream region in relation to the gene.

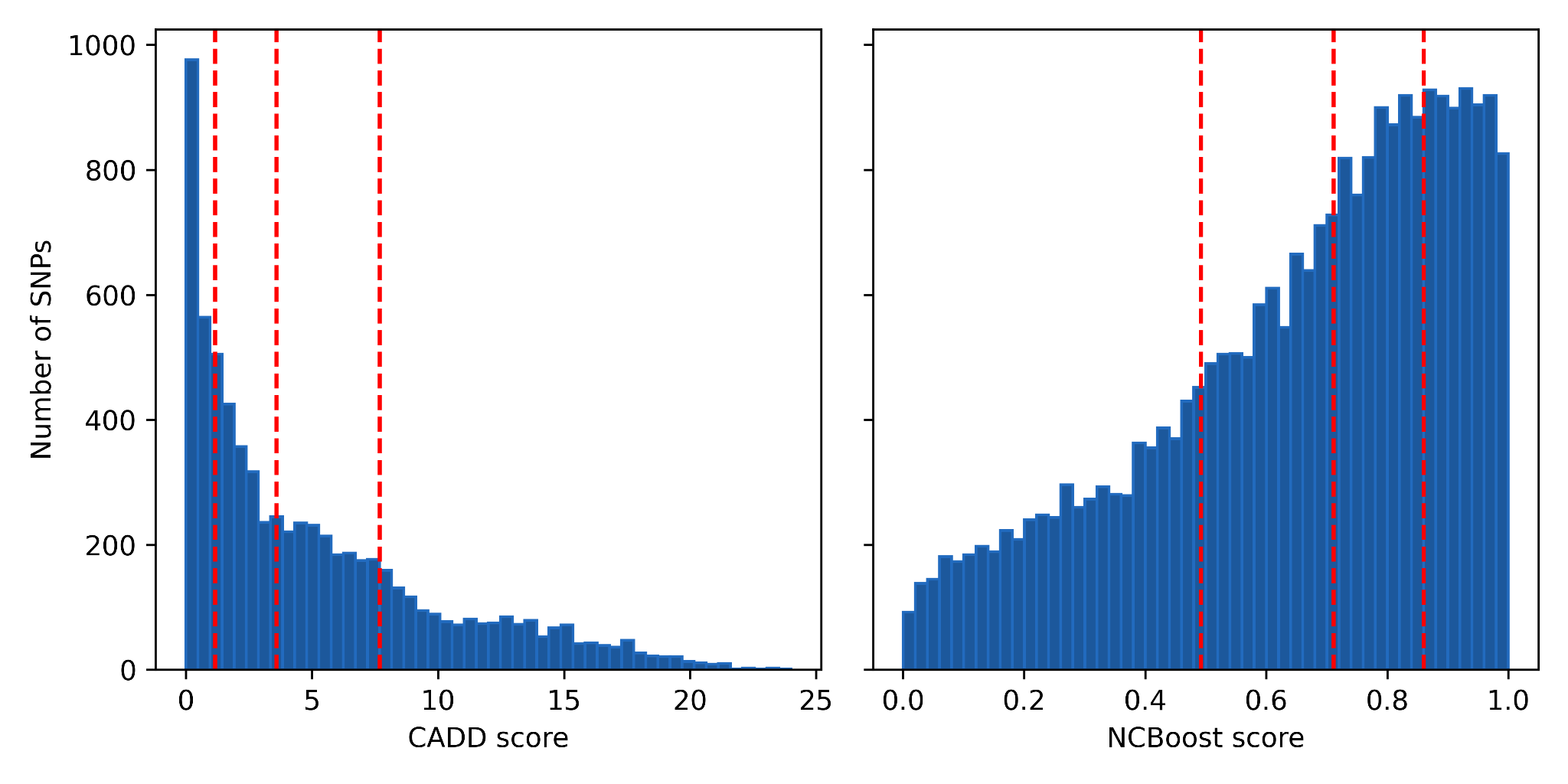

**Figure S14. Distribution of CADD and NCBoost scores of SNPs occurring within predicted functional non-coding windows in the cheetah**. Red dashed lines show the 25th, 50th and 75th percentile (from left to right) of the data. The majority of SNPs had a low CADD score, with few highly pathogenic variants, whereas the distribution of NCBoost scores is skewed towards a higher score.

Supplementary tables

Supplementary tables can be found in a separate Excel file. Table legends are displayed below.

**Table S1. Mouse & Human ATAC-seq samples.** Mouse and human ATAC-seq samples used in this study, downloaded from ENCODE (https://www.encodeproject.org/). For each sample, the species, tissue, age, sex, health status and ethnicity are provided (where relevant). Experiment, library and read accessions are provided alongside the relevant DOI. For this study, raw ATAC-seq read files (fastq) were used, but information for all files related to each sample is provided.

**Table S2. Dog ATAC-seq samples.** Dog ATAC-seq samples used in this study, downloaded from BarkBase (https://www.barkbase.org/). For each sample, the species, tissue, individual and corresponding accessions are provided.

**Table S3. Reference genomes for Felidae alignment.** Reference genomes used to generate an alignment of all chromosome-level felid genomes, which was generated with Progressive Cactus and used to calculate conservation scores.

**Table S4. Genes with previously reported deleterious coding mutations.** Genes previously identified to have deleterious mutations within protein-coding exons. The gene name and ID (where provided) are listed alongside the study of origin and whether the gene is also found in other studies (see Duplicated). Any additional information about the mutations previously identified is also listed, such as a premature termination codon (PTC) or the impact of the variant. Alongside this, whether the gene has an associated non-coding deleterious variant in this study is marked as Y (yes) or N (no). The number of unique variants associated with each gene is listed as well as the number of individuals carrying these variants. Gene lists were sourced from five studies: Dobrynin et al. (2015) Genome Biology https://doi.org/10.1186/s13059-015-0837-4; Samaha et al. (2021) BMC Genomics https://doi.org/10.1186/s12864-021-07899-2; Meißner et al. (2024) Scientific Reports https://doi.org/10.1038/s41598-024-54076-y; Peers et al. (2025) Evolution https://doi.org/10.1093/evolut/qpaf005; Peers et al. (2026) Molecular Ecology https://doi.org/10.1111/mec.70513. Genes with no gene name (marked with NA) are greyed out, alongside duplicated entries.

**Table S5. Mann–Whitney U significance values.** Significance values for pairwise Mann–Whitney U tests, run using SciPy Stats (Virtanen et al. 2020). Three sets of tests were run (ExplaiNN score, Log-transformed distance to nearest gene and Median PhyloP score) for each pairwise comparison between the four classes: true positives (TP), false positives (FP), true negatives (TN), and false negatives (FN).

**Table S6. TF motifs learned by ExplaiNN.** Transcription factor (TF) motifs learned by ExplaiNN for each species-specific model (Models 2, 3, 4). The importance for each filter for predicting positives (ATAC-seq peaks) and negatives (non-ATAC-seq peaks) is shown alongside the corresponding JASPAR motif and the significance of the JASPAR motif predictions.

**Table S7. Transcription factor binding motifs impacted by putatively deleterious non-coding SNPs.** The number of putatively deleterious non-coding SNPs overlapping with each transcription factor binding motif. Motifs were identified using the Jaspar 2024 Vertebrate Core Non-redundant motif database (https://jaspar.elixir.no/downloads/) with FIMO (MEME suite v5.1.0) (Grant et al., 2011).
